# Targeting of Mammalian Glycans Enhances Phage Predation in the Gastrointestinal Tract

**DOI:** 10.1101/2020.07.20.212829

**Authors:** Sabrina I. Green, Carmen Gu Liu, Xue Yu, Shelley Gibson, Wilhem Salmen, Anubama Rajan, Hannah E. Carter, Justin R. Clark, Xuezheng Song, Robert F. Ramig, Barbara W. Trautner, Heidi B. Kaplan, Anthony W. Maresso

**Author notes:** Address correspondence to Anthony Maresso,.

## Abstract

The human mucosal surface consists of a eukaryotic epithelium, a prokaryotic microbiota, and a carbohydrate-rich interface that separates them. Bacteriophage parasitize the prokaryotes but are not known to associate with eukaryotic cells. In the gastrointestinal tract, the interaction of these two domains influences the health of the host, especially colonization with invasive pathobionts. Antibiotics may be used but they also kill protective commensals and lack the physio-chemical properties to be specifically and optimally active in this complex milieu. Here, we report a novel phage whose lytic cycle is enhanced in intestinal environments. The enhanced activity is encoded in its tail fiber gene, whose protein product binds human heparan sulfated proteoglycans and localizes the phage to the epithelial cell surface, thereby positioning it near its bacterial host, a type of locational targeting mechanism. This finding offers the prospect of developing epithelial-targeting phage to selectively remove invasive pathobiont species from mucosal surfaces.

**Graphical Abstract:** 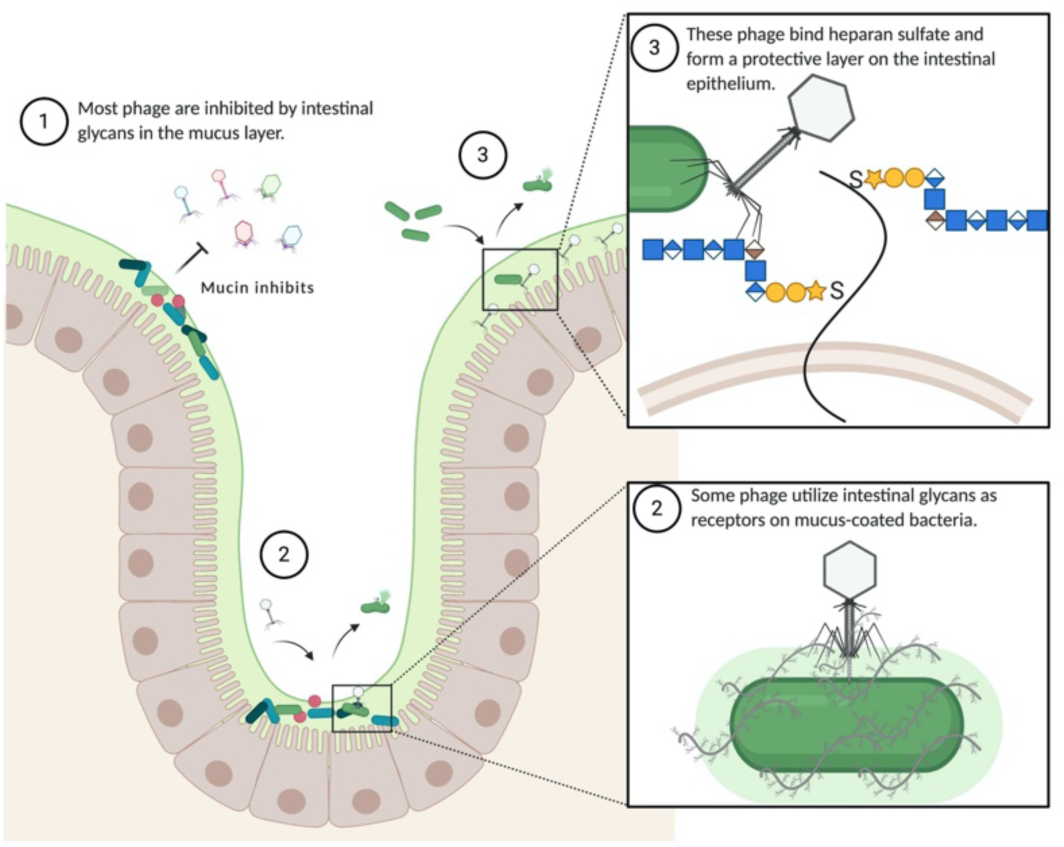

Model showing (1) mucins from the intestinal mucus layer inhibit phage infection, (2) phage ES17 can bind to mucin and utilize other intestinal glycans as a receptor to infect and kill mucus-coated bacteria, and (3) phages like ES17 can be utilized to coat the intestinal epithelium by binding heparan sulfate glycans to protect from invasive pathogen infection.

## Introduction

The human mucosal surface is a diverse community comprised of bacteria, viruses, fungi, and human epithelial, immune, and stem cells. The chemistry of the surface is complex—commonly composed of proteins, lipids, nucleic acid, and small molecule metabolites (Sansonetti, 2004; Nataro, 2005). The pH, ionic environment, and physio-mechano properties also contribute to this dynamic surface. All of these factors influence the biology, and, in turn, disease and treatment. Research over the last decade has revealed the importance of mucosal homeostasis. Invasive pathobionts may colonize, grow, and cause acute or chronic infections, some of which may become systemic and life-threatening (Johnson JR, 2002; Russo and Johnson, 2003). The bacterial composition of this intestinal community varies in association with human illnesses such as cancer, diabetes, neurological illness, obesity, and cardiovascular disease (De Martel *et al*., 2012; Sampson and Mazmanian, 2015; Maruvada *et al*., 2017; Vatanen *et al*., 2018; Tang, Li and Hazen, 2019).

Antibiotics have revolutionized medicine. The meteoric rise of multidrug resistance threatens a returns to pre-antibiotic days by 2050 (O’Neill, 2016). In addition, it is increasingly recognized that antibiotics act as a commensal destroying napalm. The antibiotics developed commercially have focused on inhibition of key processes common to all bacterial species, for example, the inhibition of protein or cell wall biosynthesis. Current antibiotics lack the specificity to selectively target a causative pathobiont, which in the mucosal environment, might be the major driver of the disease among the backdrop of hundreds of benign, even beneficial, symbionts. Moreover, the core chemical backbones of antibiotics (e.g. *β*-lactam antibiotics) somewhat limit the ability to develop new modified versions. Furthermore, these modifications improve enzymatic activity (e.g. increase affinity or induced suicide inhibition) but has not enhanced the drug’s activity in the human mucosal environment. Mucosal-active, targeted drugs that specifically control the bacterial composition of this complex environment without perturbing ecological balance would promote the health of the nasopharyngeal, lower respiratory, gastrointestinal, and urogenital epithelium.

The rise of multidrug resistance has precipitated the search for alternative antibacterial approaches, including the use of bacteriophages (phages) which are bacterial viruses used to treat infections. Although phage therapy has been used in Eastern Europe for decades, only in the past few years has it been employed for compassionate use cases in the U.S. and Europe and in clinical trials (Schooley *et al*., 2017; Dedrick *et al*., 2019; Jault *et al*., 2019). In theory, phage offer numerous advantages over antibiotics, including that they can be specific towards a given species of bacteria resulting in potential microbiome sparing, “generally regarded as safe” or GRAS attributes, allowing them to be given in high doses, and diverse, as phage mixtures can be used to either broaden the range or reduce the frequency of resistance or both (Loc-Carrillo and Abedon, 2011; Grose and Casjens, 2014; Moye, Woolston and Sulakvelidze, 2018). However, their greatest attribute has been speculated to be their abundance – estimated to be 10^31^ on Planet Earth. The phage genosphere—all the collection of genes and their unique biology—sometimes referred to as “viral dark matter”, may encode novel features that allow for enhanced lytic activity towards bacterial pathogens (Youle, Haynes and Rohwer, 2012; Brum *et al*., 2016; Terwilliger *et al*., 2020).

Reasoning that this genosphere may include phage with unique phenotypes that facilitate lytic activity against pathobionts at mucosal environments, we report here a novel podovirus of the genus *Kuravirus* that infects pathogenic *E. coli* using positional targeting to human heparan-sulfated proteoglycans.

## Results

### The gastrointestinal tract is prohibitive to phage therapy

*Escherichia coli* are gram-negative bacilli found in a variety of environments, including the human intestinal microbiome, with > 90% of people colonized (Mitsuoka, Hayakawa and Kimura, 1975). Although most strains are benign, *E. coli* is particularly troubling because of its propensity to undergo horizontal transfer of genes encoding virulence factors and antimicrobial resistance (Huang *et al*., 2001; Antão, Wieler and Ewers, 2009; Price *et al*., 2013; Mathers, Peirano and Pitout, 2015; Poole *et al*., 2017). Thus, several pathotypes have arisen that are associated with human disease. InPEC (Intestinal <u>p</u>athogenic *<u>E</u>. coli*) are associated with diarrhea and gastroenteritis. ExPEC (<u>E</u>xtraintestinal <u>p</u>athogenic *<u>E</u>. coli*) are associated with systemic infections of the urinary tract, brain, peritoneum, peripheral organs, blood, and in-dwelling devices, resulting in 9 million infections per year (Russo and Johnson, 2003; Colpan *et al*., 2013; Poolman and Wacker, 2016). The pandemic ExPEC sequence type 131 or ST131 possess a rare combination of multidrug resistance and enhanced virulence (Pitout, 2012; Johnson *et al*., 2013; Banerjee and Johnson, 2014). These strains readily colonize the human intestine, which can become a reservoir, prior to extraintestinal infection (Nicolas-Chanoine, Bertrand and Madec, 2014). Previous studies reported that phage HP3, a lytic myovirus isolated from environmental reservoirs of ExPEC, could reduce ST131 bacteremia and disease severity in murine models of infection (Green *et al*., 2017a; Ma *et al*., 2018). Since the human gastrointestinal tract is the primary reservoir of ExPEC ST131, we wondered if phage HP3 could act prophylactically to reduce or eliminate ExPEC burden in the intestine. To test this, mice were orally gavaged with an ExPEC ST131 clinical isolate then treated with phage or an antibiotic as illustrated in Figure 1A. Untreated mice sustained stable bacterial colonization during the course of the experiment (6 days) (Figure 1B). When phage HP3 were given to animals via water or a daily gavage, the levels of ExPEC were indistinguishable from the untreated control at the end of the experiment. However, no ExPEC was detected at any time point in the antibiotic-treated group. Interestingly, phage HP3 was detected, and active, in the stool of phage-treated mice, even on day 4, indicating the lack of ExPEC reduction was not due to a lack of delivery to the intestinal environment or to inactivation of the phage (Figure 1C). It should also be noted that despite having higher levels of phage upon gavage during days 1-4, the phage was no more effective at removing ExPEC than phage given in the water, suggesting that in this experiment there was no dose dependent effect of phage. In addition, as much as 10^5^ (PFU/g) phage were found in the murine intestinal tissue (including cecum and colon) on day 6, the final day of the study, yet there was little to no clearing of ExPEC compared to the untreated control in these tissues (Figures 1D and 1E). Importantly, antibiotic treatment significantly reduced the number of operational taxonomic units (OTUs) and diversity, as determined by the Shannon diversity index, which takes into account species richness and distribution (Figure 1Fi, ii, p = 0.007, p = 0.009). Also, a Principle component analysis (PCA) of beta diversity demonstrated that mice in the antibiotic cohort clustered together and away from the untreated and phage groups, suggesting antibiotics had a more profound effect on the microbiome than did phage (Figure 1Fiii). Finally, we assessed phage killing in a modified “cecal medium” or CM that is derived from the murine cecal contents and designed to simulate the luminal complexity of the mammalian intestine (Figure 1G).

**Figure 1:**
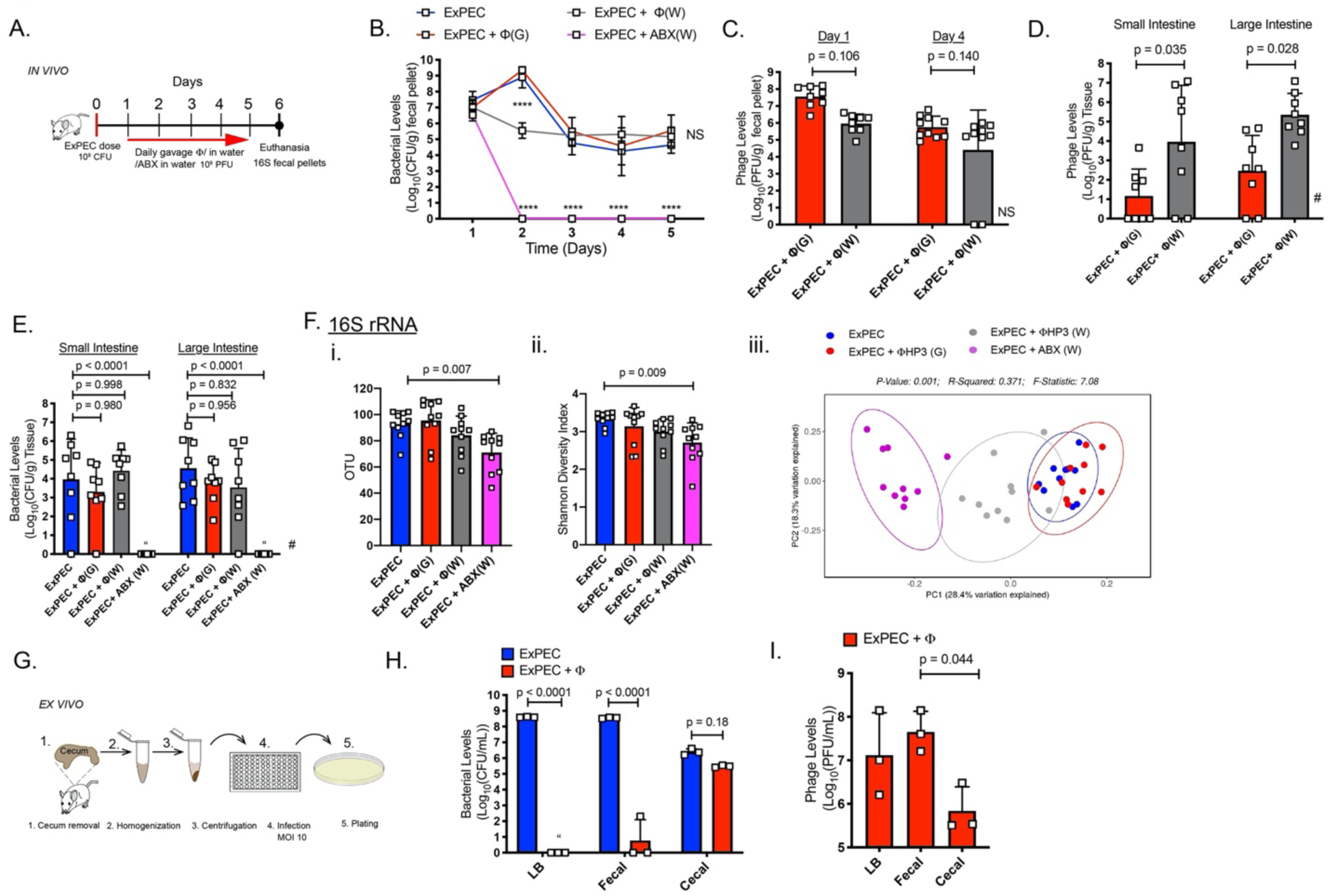
The gastrointestinal tract is prohibitive to phage therapy. (A) Mouse colonization diagram. (B) Intestinal (fecal) colonization of ExPEC (C) and phage HP3 levels. (D) Day 6 intestinal tissue HP3 levels and (E) ExPEC levels. (Fi) OTU from 16S rRNA analysis of fecal pellets day 6. (Fii) Shannon diversity values (Fiii) PCA of beta diversity. Symbols (squares) indicate individual mice, except in (B) squares indicate mean. Exp. performed 1X. N=8-10 where N indicates individual mice. **** p < 0.0001, NS not significant,“ = none detected, # missing values due to histological analysis of 2 mice/group. Mean, ±SD shown. (G) Cecal medium (CM) development. (H) ExPEC levels after 4.5 hours growth and (I) phage HP3 levels. Experiment performed 1X. Squares (N=3) indicate individual biological replicates from independent cultures. Mean, ±SD shown.

Whereas phage HP3 completely abolished ExPEC in LB (nearly a 9-log drop in levels and no detectable live bacteria), and nearly abolished it in a slurry of fecal pellets taken from the same murine host (∼ 8-log drop in levels), there was little to no phage-based killing in CM, despite recovering nearly 10^6^-10^7.5^ PFU/mL (Figures 1H and 1I) of phage. The result was the same when the experiment was repeated anaerobically (data not shown). These results mirrored those from the ExPEC colonization model and indicate there is/are factor(s) present in the mammalian GI that inhibit this lytic phage.

### The inhibitory component is mucin

We wished to understand the reasons phage HP3 was ineffective in this intestinal microenvironment. Previously, we had shown that phage HP3 utilizes calcium for efficient infection, suggesting the effect may be nutritionally-based (Ma *et al*., 2018). However, addition of calcium did not increase phage HP3 killing in CM (data not shown). We next tested whether the inhibition might be related to the presence of live bacterial microbiota in CM. However, ExPEC killing with phage was not enhanced with removal of the microbiota with a broad range of antibiotics, including inhibitors of protein synthesis, cell wall, and DNA synthesis (Figure S1i-iii – note that the antibiotics efficiently killed a commensal, antibiotic-sensitive, *E. coli* that was spiked into the CM, iv). Furthermore, the inhibitory effect was maintained when a related, and equally as effective phage EC1 was used in the same experiment, thereby indicating the inhibitory effect was not solely due something specific for phage HP3 (data not shown). Arriving at no resolution as to what the inhibitory factor may be, we decided to test more drastic treatments for their ability to restore phage killing in CM. First, the CM was heat-treated (Figure 2Ai, HT CM). Interestingly, heat treatment led to greater than a 6-log improvement in ExPEC killing by phage HP3 (Figure 2Aii, p < 0.0001). Similarly, when CM was filter-treated (Figure 2Ai, FT CM), there was no detectable levels of ExPEC in the medium after treatment with phage HP3 (Figure 2Aii), and, this observation was extended to another phage showing inhibition in CM, EC1 (Figure S2A, p < 0.0001). Thus, the inhibitory component was large (retained on 0.22 micron filter) and sensitive to boiling.

We reasoned intestinal mucins might fit this profile due to their highly associative and sticky properties (captured on a filter) and as proteins they would be sensitive to heat. Mucins are extensively glycosylated proteins found throughout the gastrointestinal system, which form a layer between the intestinal epithelial cells (IECs) and the commensal or pathogenic microbiota (Johansson and Hansson, 2016). Also, they can function as receptors for microbes (McGuckin *et al*., 2011). To test whether mucins were inhibiting bacterial killing by phage, we devised another method whereby CM was separated via high speed centrifugation into soluble (S CM) and insoluble (INS CM) forms (Figure 2Bi). We hypothesized that INS CM would contain mucin, since large intestinal mucins are normally present in this portion, and be inhibitory to phage killing, whereas the S CM would not have these properties (Carlstedt *et al*., 1993). Indeed, phage killing in unprocessed CM or INS CM was inhibited to a greater extent than in S CM (Figure 2Bii). To more directly test the hypothesis that mucin was the inhibitory factor, the mucolytic drug N-acetyl cysteine (NAC) was added to INS CM and porcine gastric mucin (1.5 % m/v) added to S CM (Figure 2Bi-ii). Indeed, a 5-log reduction in phage killing of ExPEC was observed in INS CM upon addition of NAC, which also improved killing to that seen in S CM (p < 0.0001). Perhaps more compelling, the addition of mucin to S CM abrogated bacterial killing by phage to levels originally observed in CM alone (not significant p = 0.970). These results indicate that the inhibitor of phage killing in cecal medium is intestinal mucin.

Along these lines*, E. coli* is known to use mucins as a source of carbon (Chang *et al*., 2004). We reasoned that a murine host colonized with ExPEC may see a bloom upon NAC treatment due to the drug liberating the mucins for bacterial consumption and thus serve as a system to test if the reduction in aggregated mucin would promote phage HP3’s ability to kill ExPEC. Indeed, upon treatment with NAC for two weeks, ExPEC levels in the small intestine were increased and phage HP3 reduced ExPEC levels, 2.5 logs, in the small intestine (Figure S2Bi-ii). A similar trend was observed in the large intestine, but it was not substantial (<1 log reduction), though in both organ sections the data were not significant (Figure S2Biii). The less pronounced effect in the large intestine may be due to the thickness of mucus, and thus the lower likelihood for NAC to be effective at breaking up this mucus. Also, NAC’s known to be rapidly absorbed in the small intestinal tissue, thereby losing its effect in the more distal large intestine (Aruoma *et al*., 1989; Tsikas *et al*., 1998).

### Discovery of a mucin-enhanced phage

An estimated 10^31^ phages exist in nature. And the human gastrointestinal tract contains a large diversity consisting of >10^10^ phages many of which are just beginning to be discovered (Shkoporov and Hill, 2019; Sausset *et al*., 2020). Reasoning that human sewage or the feces of animals may contain phage that have evolved to target their host in high mucin environments, such as the intestinal tract, we screened our phage library (Gibson *et al*., 2019) and other phages (Table 1) recently isolated from these environments for enhanced activity in LB containing 1.5% mucin (Figure 2C). This is the same concentration that prevented phage HP3 activity in soluble cecal medium and was shown to provide strong inhibition in LB for up to 8 hours (Figure S3A). Surprisingly, only a single phage, designated phage ES17, significantly killed bacteria in the LB mucin medium (Figure 2Cii, p < 0.0001, about 1-log improvement). Surprisingly, phage ES17 was active in mucin despite being much less effective (> 3-logs) than phages HP3, J2W, Ult1, Shp1, or M1S, which completely killed ExPEC to undetectable levels in LB medium alone (Figure 2Ciii, LB). Consistent with these data, when the amount of mucin was varied from 0 to 2% and phages HP3 and ES17 were compared for lytic activity against ExPEC, phage HP3 was highly effective as the concentration of mucin was lowered to below 0.5%, but completely inhibited above that level (Figure S3B).

**Figure 2:**
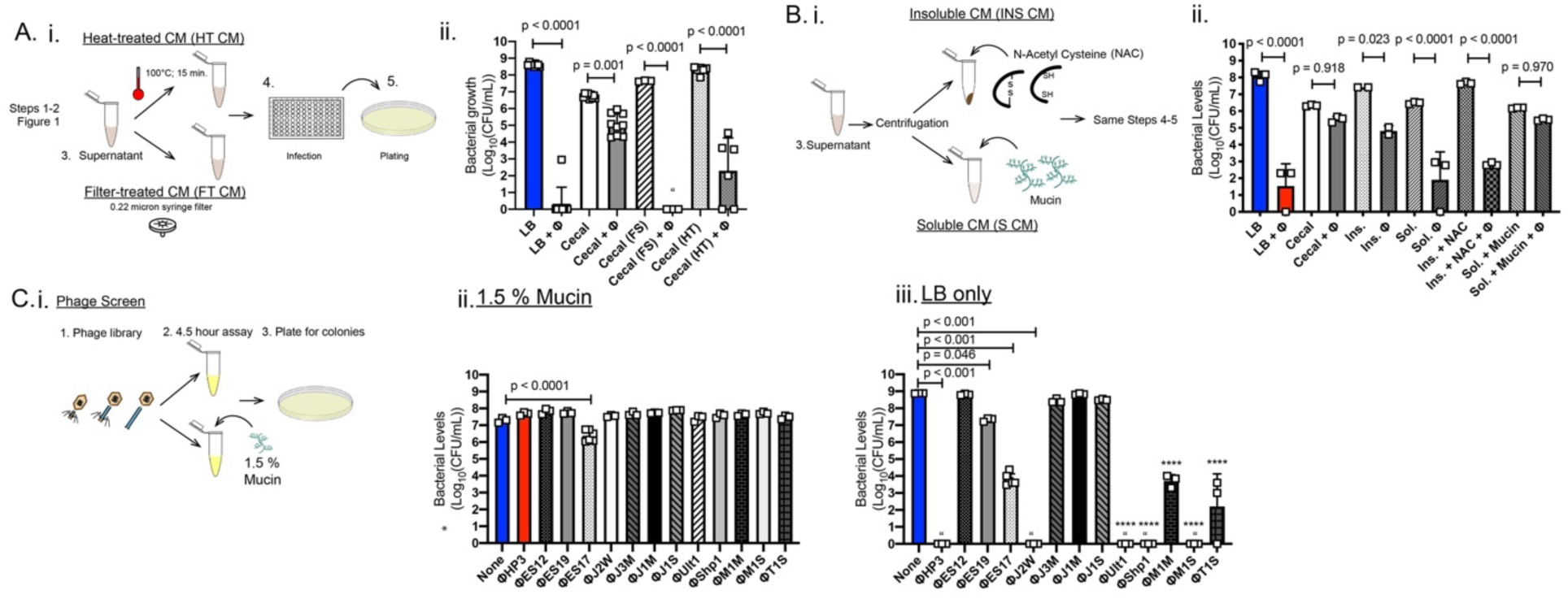
The inhibitory component is mucin. (Ai) CM was heat-treated (HT, 100°C) or filter treated (FT) prior to assay. (Aii) ExPEC levels in HT CM and FT CM. (Bi) CM was separated into pellet (insoluble), homogenized and treated with N-acetyl cysteine (NAC), or into supernatant (soluble) and mucin added. Following assay modified CM was plated for (Bii) ExPEC levels. (Ci) A screen of phage killing in mucin was performed using the phage in Table 1. (Cii) ExPEC levels in LB + 1.5% mucin. (Ciii) LB alone. N=3-6, **** p < 0.0001, “ = none detected. Exp. performed 1X (B), 2X (A and C). Squares indicate individual biological replicates from independent cultures. Mean, ±SD shown.

**Table 1.**
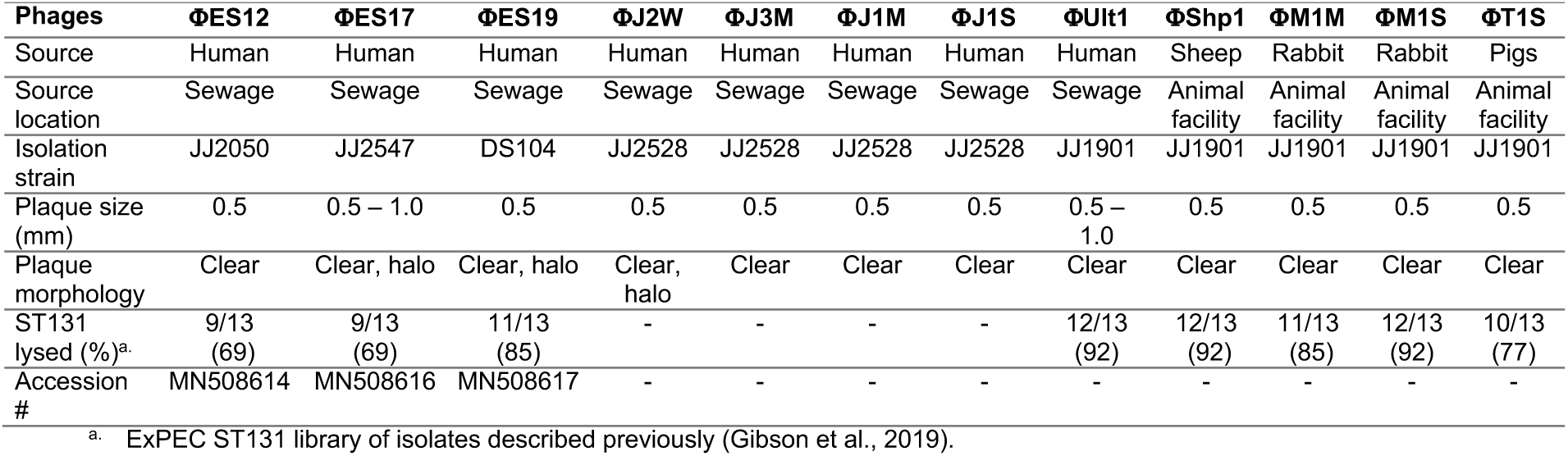
Properties of E. coli phages used in mucin screen.

| Phages | ΦES12 | ΦES17 | ΦES19 | ΦJ2W | ΦJ3M | ΦJ1M | ΦJ1S | ΦUlt1 | ΦShp1 | ΦM1M | ΦM1S | ΦT1S |
| --- | --- | --- | --- | --- | --- | --- | --- | --- | --- | --- | --- | --- |
| Source | Human | Human | Human | Human | Human | Human | Human | Human | Sheep | Rabbit | Rabbit | Pigs |
| Source location | Sewage | Sewage | Sewage | Sewage | Sewage | Sewage | Sewage | Sewage | Animal facility | Animal facility | Animal facility | Animal facility |
| Isolation strain | JJ2050 | JJ2547 | DS104 | JJ2528 | JJ2528 | JJ2528 | JJ2528 | JJ1901 | JJ1901 | JJ1901 | JJ1901 | JJ1901 |
| Plaque size (mm) | 0.5 | 0.5 – 1.0 | 0.5 | 0.5 | 0.5 | 0.5 | 0.5 | 0.5 – 1.0 | 0.5 | 0.5 | 0.5 | 0.5 |
| Plaque morphology | Clear | Clear, halo | Clear, halo | Clear, halo | Clear | Clear | Clear | Clear | Clear | Clear | Clear | Clear |
| ST131 lysed (%)<sup>a</sup> | 9/13 (69) | 9/13 (69) | 11/13 (85) | - | - | - | - | 12/13 (92) | 12/13 (92) | 11/13 (85) | 12/13 (92) | 10/13 (77) |
| Accession # | MN508614 | MN508616 | MN508617 | - | - | - | - | - | - | - | - | - |

<sup>a</sup>. ExPEC ST131 library of isolates described previously (Gibson et al., 2019).

However, phage ES17 was most effective at concentrations in which HP3 was inactive (0.5 – 1% mucin) and least active at concentrations below or above this range. Taken together, these data suggests that phage ES17 harbors unique properties that facilitate its ability to efficiently be lytic in the presence of mucin.

### Phage ES17 is a rare C3 type phage whose activity is enhanced by mucin

We sought to understand the molecular mechanism of phage ES17’s enhanced ability to find and lyse its bacterial host in mucin. Phage ES17 was determined to be a dsDNA virus of the order *Caudovirales*, family *Podoviridae* and genus *Kuravirus*. The genus *Kuravirus* was named for the River Kura in Tbilisi, Georgia where the original phage of this genus, PhiEco32, which shows close genetic similarity to ES17, was first isolated (Figure S4A) (Savalia *et al*., 2008). Kuravirus phages have similar morphological characteristics (Ren *et al*., 2019), including an elongated C3 type capsid, which is considered a rare morphotype for phages (only 13% of tailed phages have elongated heads) and short tail fibers (Ren *et al*., 2019). Electron microscopy confirmed ES17 to have these same morphological characteristics (capsid >100 nm – Figure 3A). Similar to other phages of this genus, ES17 has a small genome size consisting of 75,007 bps with 123 predicted ORFs (Figure 3B).

**Figure 3:**
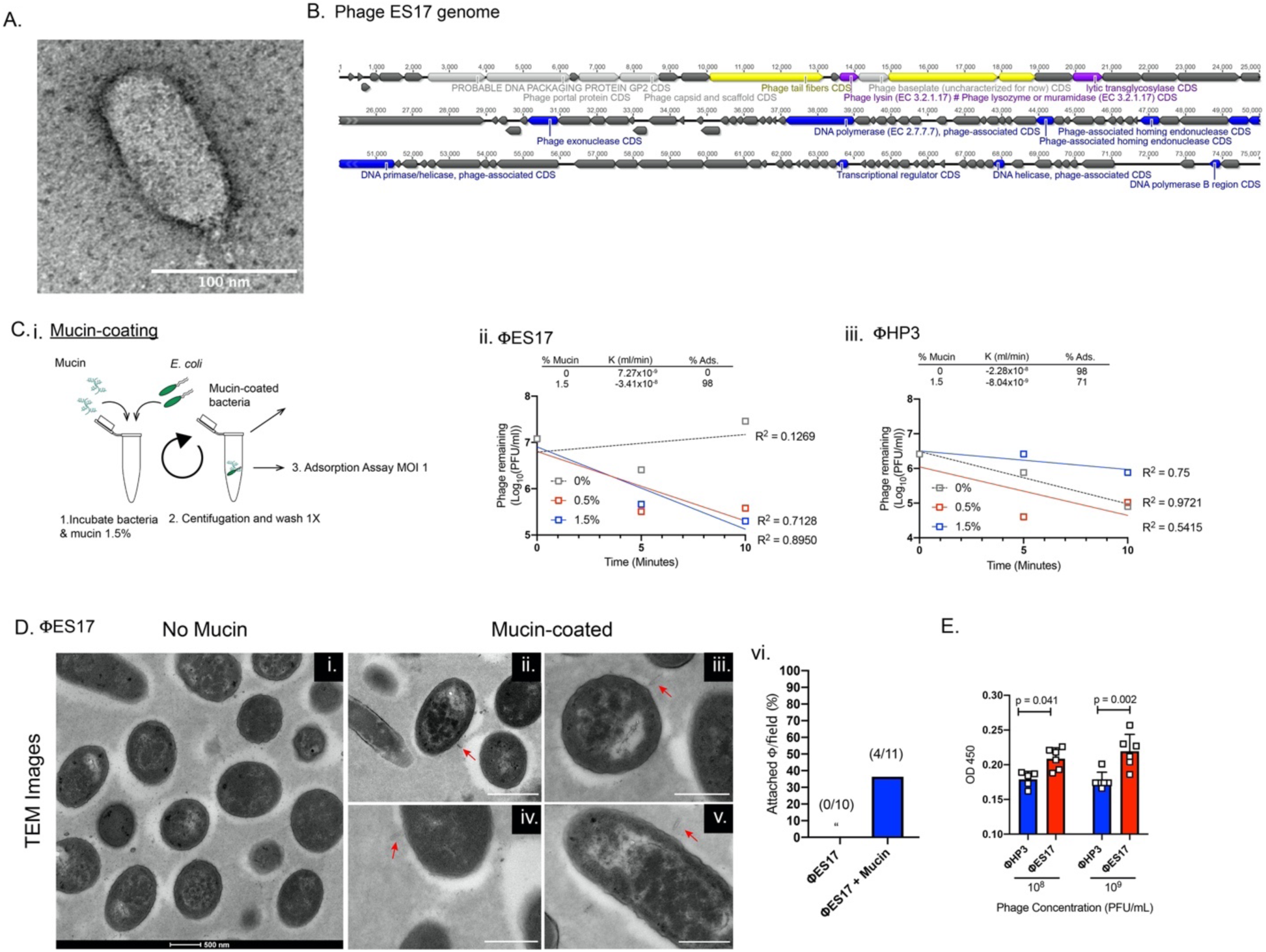
Phage ES17 is a rare C3 type phage with avidity for mucin. (A) TEM of ES17. (B) Genome of ES17. (Ci) ExPEC was incubated in 0%, 0.5% and 1.5% mucin, centrifuged, washed and phage added for an adsorption assay. (Cii) Phages ES17 and (Ciii) HP3 adsorption. Adsorption constant (K) and % adsorbed in 10 min indicated. Regression line generated from mean values of phage remaining for time points indicated. (Di-v) TEM of phage ES17 and ExPEC fixed after 10 min. adsorption. White bar is 500 nm. Red arrows indicate phage (Dvi) Attached phages per field view. (E) Mucin-ELISA results for phages HP3 and ES17. Squares indicate individual biological replicates from independent cultures. Exp. performed 1X (C) and (D), 2X (E). N=6, “ = none detected. Mean, ±SD shown.

In order to determine why this phage is distinct from other phages that lack activity in mucin-rich environments, we examined the ability of phages ES17 and HP3 to adsorb to their *E. coli* hosts. Previous data had shown that 98% of HP3 was adsorbed in 10 minutes, whereas only 32% of ES17 was adsorbed in that time (Gibson *et al*., 2019). We wondered whether the addition of mucin could improve phage ES17 adsorption and inhibit the adsorption of phage HP3. A modified adsorption assay was utilized for this experiment using ExPEC “coated” with mucin. Briefly, the bacteria were incubated in different concentrations of mucin (0%-1.5%), pelleted, and washed to remove any mucin that did not adhere to the bacterial surface (Figure 3Ci). Next, a standard adsorption assay was conducted with phages ES17 and HP3. Interestingly, phage ES17 showed no adsorption to ExPEC in 10 minutes in the absence of mucin; however, if first incubated with 1.5% mucin, adsorption increased to 98% (Figure 3Cii). In contrast, 98% of phage HP3 was adsorbed without mucin and the adsorption dropped to 75% in its presence (Figure 3Ciii). Transmission electron microscopy (TEM) analysis of these samples at 10 minutes demonstrated that only in the presence of mucin was phage ES17 found bound to the bacterial surface (red arrows) (Fig 3Di-vi, 4 out of 11 images of ExPEC + mucin had visible evidence of ES17 compared to 0 out of 10 in the absence of mucin). TEM imaging of phage HP3 adsorption showed the presence of bound phage in images with and without mucin present (red arrows) (Figure S4Bi-ii). In this case, 5/14 of the imaged fields showed phage HP3 bound with mucin-coated ExPEC compared with 7/14 bound without mucin (Figure S4Biii). Finally, we adapted an ELISA-like approach to determine if phage ES17 preferred binding to surfaces coated with mucin, a hypothesis consistent with our data. Indeed, when mucin was bound to an ELISA plate, phage ES17 bound to the mucin-surface at higher levels than phage HP3 (Figure 3E, p = 0.041 and p = 0.002). Taken together, using three different approaches, these data obtained suggests that phage ES17 binds mucin, a property that may enhance its ability to infect *E. coli* in mucin-rich environments.

### Phage ES17 may bind bacterial polysaccharides

As indicated above, phage ES17 is a close relative of the kuravirus phiEco32 and annotated as the same phage type. ES17’s putative tail fiber protein (ES17-TFP) showed high homology (64% identical) to a tail fiber protein in another lytic podophage, the T7-like bacteriophage LM33_P1, which also targets ST131 strains (Dufour *et al*., 2016). This homology is in the C-terminal receptor binding region (Figure S5Ai). The N-terminal region is similar to vB_EcoP_WFI101126 (47% identical), a phage with C3 like morphology similar to phage ES17 (Figure S5Aii)(Korf *et al*., 2019). The T7-like phage tail fibers have recently been characterized and shown to have knob-like tips, which we also observed in the EM images of phage ES17 (Figure 3A) (Garcia-Doval and van Raaij, 2012). T7-like phage tail fibers have been shown to possess endosialidases that target surface sugars, such as capsule-forming polysaccharides (Garcia-Doval and van Raaij, 2012). A BLAST analysis revealed that ES17-TFP contains a putative pectinesterase (E value 7.45e-03, 369 bp) with high homology to phage lyases (Figure 4A). Modeling of the predicted structure of ES17-TFP indicated it may contain a pectin lyase fold (Figure S5B, IPR011050). These lyases cleave bacterial polysaccharides and are found in biofilm-degrading phages (Latka *et al*., 2017). Additionally, this enzyme was not found in phage HP3. Thus, we formed the hypothesis that ES17 might degrade and infect biofilms.

**Figure 4:**
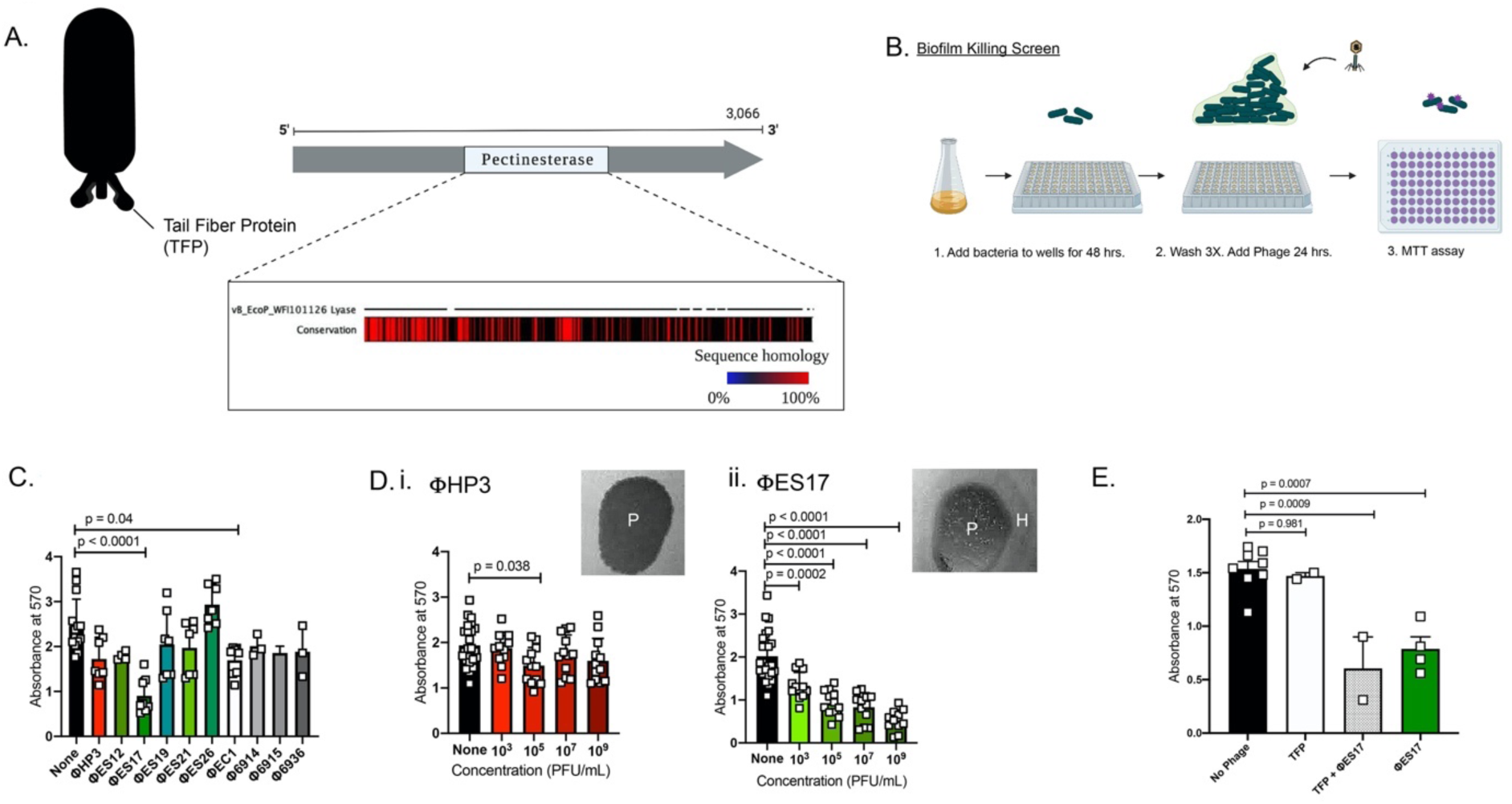
Phage ES17 may bind bacterial polysaccharides. (A) ES17 TFP domain compared to vB_EcoP_WFI101126 lyase. (B) DS515 was grown o/n in TSB and inoculated on a plate for 48-hr biofilm growth. A phage library was screened with MTT used to quantitate biofilm forming bacteria. (C) DS515 levels with phage library (N=3-8). (Di) DS515 levels with HP3 and (Dii) ES17 (N=12). Plaques “P” from phages on pictured with or without halo “H” (E) ExPEC levels with purified ES17-TFP in CM assay (N=2-8). Squares indicate individual biological replicates from independent cultures. Exp. performed 2X. Mean, ±SD shown. Figure created with BioRender.com.

ExPEC is a primary cause of catheter-associated UTI (CAUTI), often associated with the formation of biofilms on the catheter (Russo and Johnson, 2003). For this study, we used a clinical isolate (DS515) from a CAUTI patient, which forms significant biofilms *in vitro* (data not shown). We tested the effect of various *E. coli* phages on DS515 biofilms using a colorimetric assay with a metabolic substrate, MTT, to determine the sensitivity of the biofilm bacteria to the phage (Figure 4B). As determined in Figure 4C, of the 10 *E. coli* phages assessed, which were shown to be lytic towards the clinical isolate, only two phages significantly educed *E. coli* viability after establishment of the biofilm, and only one of these three, ES17, reduced the biofilm by more than half. Since phage HP3 showed moderate levels of killing of *E. coli* in biofilms, we compared its ability to phage ES17 over a 6-log titration range. Strikingly, as little as 10^3^ PFU/ml of phage ES17 statistically reduced biofilm viability, which was greater than that achieved with 10^9^ PFU/ml of phage HP3 (Figures 4Di-ii). In addition, phage ES17 showed a dose dependent response at each concentration with the greatest effect seen at 10^9^ PFU/ml (p < 0.0001) (Figure 4Dii). Also, we found that phage ES17 formed large plaques with halos on DS515 and related clinical isolates (picture above Figure 4Dii). This is in contrast to phage HP3 which showed no halo formation (picture above Figure 4Di). This halo formation is indicative of EPS (extracellular polymetric substance)-degrading activity (Knecht, Veljkovic and Fieseler, 2020).

We reasoned ES17-TFP may be responsible for the cleavage of the EPS in the biofilm, thereby allowing phage ES17 to gain access to *E. coli* in the biofilm. Thus, we cloned the DNA encoding ES17-TFP into an expression vector and purified it from *E. coli*. A highly induced band of the expected molecular weight of ES17-TFP (107 KDa) was observed upon SDS-PAGE separation of the eluted fractions (Figure S5C). This band was not observed in the vector only control elutions (data not shown). When ES17-TFP was added to the biofilms, no change in bacterial viability was observed, nor did it enhance ES17 killing of *E. coli* in biofilms when added separately (Figure 4E). Likewise, when incubated in CM, or examined for its ability to degrade mucus using a gel-shift assay in an anti-mucin Western blot, it did not enhance ES17 activity or degrade mucus (data not shown). Furthermore, phage ES17 by itself also did not demonstrate this activity. These data suggests that phage ES17’s unique killing abilities are not dependent on degradation of polysaccharides (e.g. mucin or bacterial). This leaves open the question of how this tail fiber protein may enhance killing in carbohydrate-rich environments.

### Phage ES17 binds human heparan sulfated proteoglycans

Phage ES17 harbors an enhanced ability relative to other *E. coli* phages to find its bacterial host in environments in which carbohydrates are a prominent chemical component (examples from above include cecal medium, mucin-rich broth, and biofilms). A structural analysis of modeled ES17-TFP showed a high similarity to a phage K5 lyase binding domain (E-value=4E-12). The predicted structure of ES17-TFP (blue) and K5 lyase are pictured in Figure S6Ai (PDB:2X3H; red) with identical residues colored yellow. Phage K5 binds K5 capsular polysaccharide and acts as a K5 polysaccharide lyase (Hanfling *et al*., 1996). The K5 *E. coli* capsule is made of a repeating disaccharide that is identical to the precursor of heparin and heparan sulfate (HS), a linear polysaccharide, present in glycosaminoglycans (HSPGs, heparan sulfate proteoglycans). (Figure S6Aii) These proteoglycans are found on mammalian cells and in mucus (Monzon, Casalino-Matsuda and Forteza, 2006). Also, mucins with similar structures to heparan sulfate/heparin (*α*-linked GlcNAC or N-Acetyl-D-glucosamine) are present intestinally and found in PGM (Fujita *et al*., 2011).

We reasoned that ES17’s enhanced activity might be due to a novel ability to bind mammalian polysaccharides found on glycoproteins. This would be a novel mechanism by which a phage could localize to its host, thereby explaining its enhanced activity in a mucin-rich environment and in EPS biofilms. To test this idea, we assessed the ability of purified ES17-TFP to bind to a glycan array containing over ∼ 860 unique glycan structures from porcine gastric mucin (PGM), glycosaminoglycans (GAGs) and a variety of synthetic and naturally sourced glycans generated by the Consortium for Functional Glycomics (CFG) (see supplementary methods). No or very low relative fluorescent units (RFU), a proxy for binding, was observed for a wide array of mammalian glycans, including those purified from porcine gastric mucin (Figure S6Bi-ii). However, surprisingly, there was an increase of several orders of magnitude in RFU (RFUs > 2000) observed for binding to the GAGs containing heparan sulfate (ID # 64-173), but not the structurally similar GAGs hyaluronic acid #1-20 or chondroitin sulfate #21-63 (Figure S6Bii-iii). The finding that purified ES17-TFP binds human heparan sulfated proteoglycans provides a possible mechanism to explain why phage ES17 demonstrates enhanced activity in intestinal environments.

### ES17 binds to the surface of human intestinal enteroids (HEIMs)

Human intestinal enteroids (HIEs) are organotypic, higher-order cultures that have become popular as surrogates to model the human intestine. They can be grown as 3-dimensional structures complete with a lumen and crypt/villus axis or as 2-dimensional monolayers that facilitate host-pathogen interactions (In *et al*., 2016; Saxena *et al*., 2016). These cultures are also useful because they express a variety of glycans found in the human intestine, including mucins and proteoglycans (Saxena *et al*., 2016). Human intestinal enteroid monolayers (HIEMs) were derived from colonic stem cells following differentiation for 5 days in high Wnt medium. Phage ES17 or HP3 were added to confluent HIEMs for 1 hour, extensively washed, and visualized by immunofluorescence microscopy using antibodies raised against each phage. Little to no detectable phage HP3 was observed on the HIEMs intestinal epithelial cell (IEC) surface, though antibodies generated robust signal and specificity towards the phage when HP3 was fixed on slides alone (Figure 5A and S4C). Phage ES17 (green) bound evenly to the IECs on the apical side, including areas where prominent Muc2 (red) localization was observed, but also on areas where there was no muc staining (Figure 5Aii). In order to determine whether phage ES17 bound to the IECs of HIEMs via heparan sulfate, we pre-treated the HIEMs with heparinase III to enzymatically remove HSPGs and then assessed phage binding. Indeed, HIEMs treated with heparinase significantly reduced the levels of bound ES17, both qualitatively and quantitatively (Figures 5Bi-vii, p < 0.0001). Next, we tested whether this binding to the epithelial cell surface also improved bacterial killing. Previous data had shown that ExPEC adheres less to intestinal enteroids compared to the diarrhea-causing pathogen Enteroaggregative *Escherichia coli* (EAEC) (data not shown). EAEC adheres to HIEMs robustly in an aggregative, mesh-like pattern and similar to phage ES17 utilizes heparan sulfate as a receptor to bind these human colonic cells (Rajan et al., 2020, paper *in review*)(Poole, Rajan and Maresso, 2018; Rajan *et al*., 2018). Thus, EAEC serves as an ideal *E. coli* strain to test the hypothesis that phage targeting HSPGs can increase killing. For this experiment we pre-coated HIEMs with phage ES17, washed, as detailed above, then infected with EAEC strain 042. HIEMs were fixed and stained using a Geimsa-Wright stain to visualize cells and bacteria, as previously described (Rajan *et al*., 2018). Infected HIEMs showed robust bacterial adhesion to cells with an aggregative phenotype (Figure 5Cii, red arrows). Phage coated HEIMs, however, showed significantly reduced EAEC on the surface of the organoid (Figure 5Ciii-iv, **p = 0.007). Taken together, these data suggests phage ES17 may localize to both the mucus layer and to the IEC surface by binding to HSPGs, which likely position the phage to be in the exact location needed to find its bacterial target in the intestinal microenvironment.

**Figure 5:**
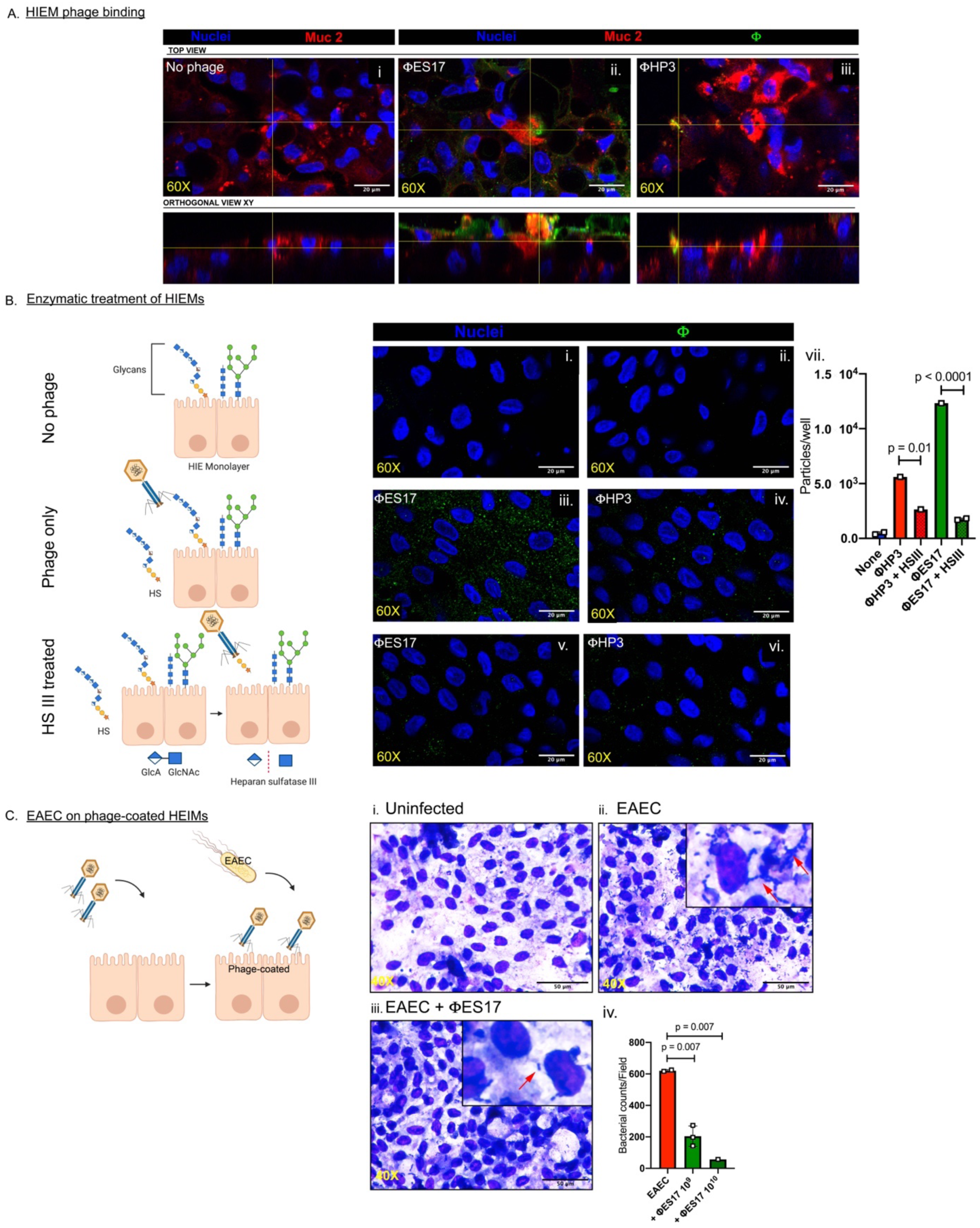
ES17 binds to the surface of human intestinal enteroids (HIEMs). HIEMs were incubated with phage for 1 hr, washed, then fixed and imaged for phage (Alexa Fluor 488, Green), intestinal cells (DAPI, Blue), and muc2 (Alexa Fluor 594; Red). (Ai) No phage added. (Aii) Phage ES17 or (Aiii) HP3 added. Monolayers pretreated with heparanase III then with phage. (Bi, ii) No phage. (Biii) Phage ES17 and (Biv) phage HP3 added. (Bv) Hep III treated HEIMs incubated with ES17 and (Bvi) HP3 (Bvii) Quantification of particles (phage) per well. (N=1-2). Images at 60X. Scale bars at 20 μm. HIEMs (Ci) uninfected or (Cii) Infected with EAEC 042 (Ciii) or pretreated with phage ES17 prior to EAEC. Images at 60X. Scale bars at 50 μm. (Civ) EAEC attached to HIEMs per field view. Mean, ±SD shown. Squares indicate individual biological replicates from independent cultures. Exp. performed 1X. Figure created with BioRender.com.

### Phage ES17 kills ExPEC in the mammalian intestine

The finding that phage ES17 demonstrated enhanced lytic activity in the presence of mucins, was the best of several screened phages in a mock luminal environment rich in mucins, and binds the human organotypic culture IECs via HSPGs, which leads to EAEC-killing on the bacterial surface, prompted an examination into whether this phage could overcome the intestinal-induced inhibition of phage lytic activity towards colonized ExPEC that was observed for phage HP3. We first tested if ES17 was effective in cecal medium. Indeed, phage ES17 showed a 2.5-log improvement in ExPEC removal in this environment compared to phage HP3 (Figure 6A, p < 0.0001). There was no effect on the number of OTUs or the Shannon diversity index in this experiment, suggesting phage ES17 was highly selective at removing only the target ExPEC strain (Figure 6Bi-ii). We next tested the effect of phage ES17 on ExPEC in a murine intestine. Animals were colonized with ExPEC as in Figure 1A and treated with either phage ES17 or HP3 (Figure 6C).The dose of phage was also increased from 10^9^ PFU to 10^10^ to also evaluate if giving more phage would improve HP3’s ability to reduce ExPEC, especially in more proximal segments as the phage slowly moves through the alimentary canal. Examination of the small and large intestine on day 6 showed phage levels were high across all groups (10^6^-10^8^ PFU/g intestinal tissue) (Figure 6D). Animals treated with phage HP3 had no detectable CFU in small intestinal tissue (Figure 6E), but had indistinguishable levels from that of the untreated control in the cecum. In contrast, every animal treated with phage ES17, except one, had no detectable levels of ExPEC in either the small or large intestine, suggesting that this lytic phage possess a unique ability to target ExPEC in complex mucosal environments such as the large intestine.

**Figure 6:**
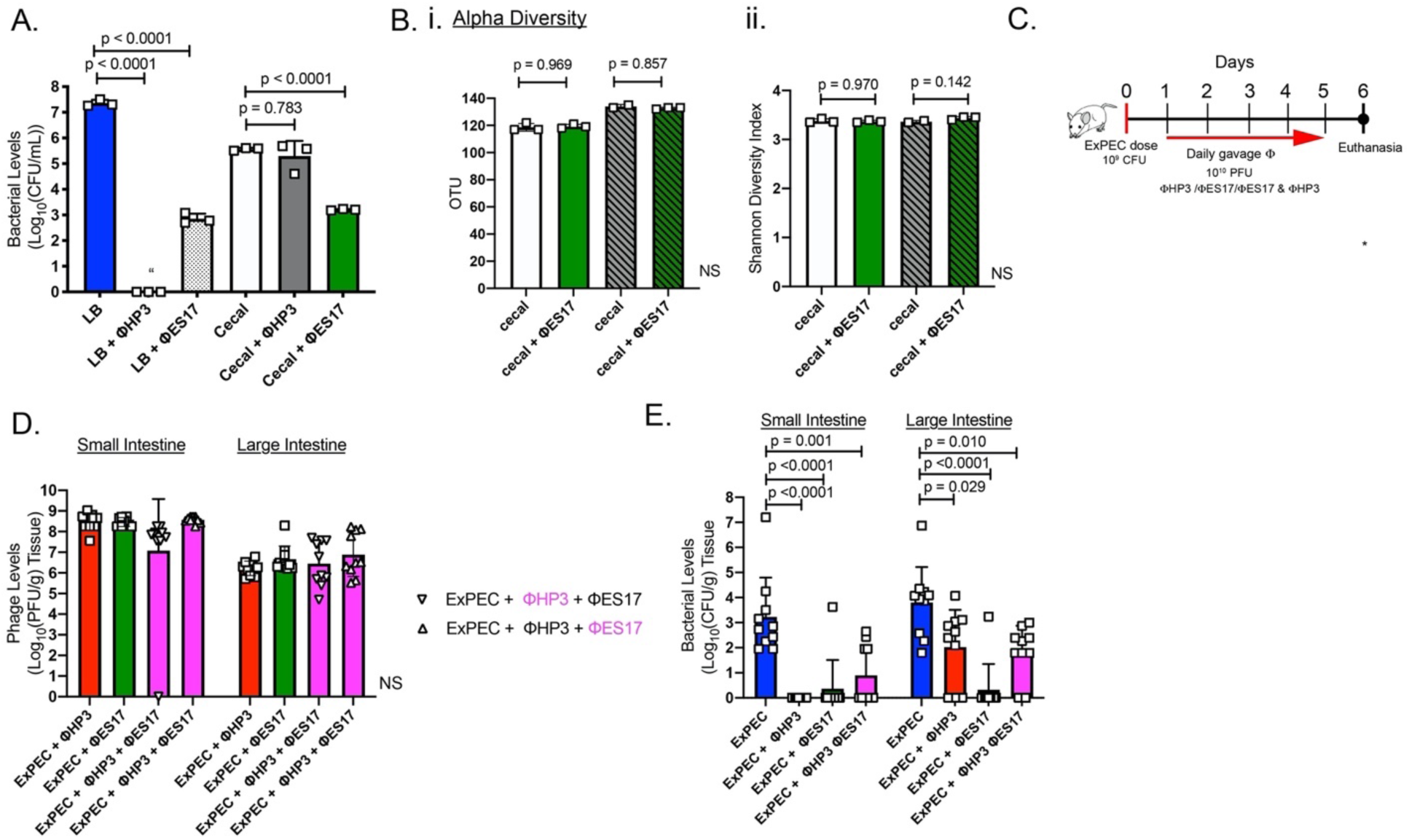
ES17 kills in the mammalian intestine. (A) ExPEC levels in CM assay with ES17 (N=3). 16S rRNA analysis of CM including (Bi) OTU and (Bii) Shannon diversity index. (C) Mice were gavaged with ExPEC then treated with no phage or gavaged daily doses of phage HP3 at 10^10^ PFU, or ΦES17 at 10^10^ PFU or both phages (2×10^10^ PFU total). (D) Intestinal (tissue) phage levels (E) Intestinal (tissue) ExPEC levels. (N=10). Symbols (squares) indicate individual mice or biological replicates derived from different cultures. Exp. performed 1X. NS not significant. “ = none detected. Mean, ±SD shown.

## Discussion

A limitation of all antibiotics is their broad killing activity and no inherent features to function well in complex human environments, including human blood, urine, or at mucosal surfaces throughout the body. In particular, the gastrointestinal tract (GIT) is a highly complex system made up of diverse organs and tissues and within that a diversity of cell types. Some of the cells and factors found at the mucosal surface may influence intestinal capacity including: intestinal epithelial cells, enteroendocrine cells, stem cells, mucus-secreting goblet cells, antimicrobial peptides (AMPs – from Paneth cells) and secretory immunoglobulin A (Mammarappallil and Elsinghorst, 2000; Johansson, Larsson and Hansson, 2011; Sato *et al*., 2011; Gribble and Reimann, 2016; Pickard *et al*., 2017). To add to this complexity, the GIT is colonized by a large number of microbes—archaea, fungi, protists and bacteria that have co-evolved with the host, termed the human gut microbiota (Pickard *et al*., 2017; Heintz-Buschart and Wilmes, 2018). A bacteriophage would have to navigate through this complex ecosystem to find and infect its host. Since there are estimated to be >10^10^ PFU/g of phages already present in the GIT, this suggests that phages can thrive in this environment (Hoyles *et al*., 2014; Shkoporov *et al*., 2018). Here, we report a novel anti-bacterial targeting strategy that seems to have evolved to position a lytic bacteriophage in the exact niche as its bacterial host. Our results indicate that; (i) phage HP3 that is lytic towards *E. coli* ST131 *in vitro* and is very effective at eliminating bacteremia in a murine model of sepsis, is ineffective when tested for this same property in the murine intestinal tract; (ii) that the inhibition is due to intestinal mucins; (iii) a medium designed to mimic the luminal environment can be used to identify phage with enhanced lytic activity in such an environment; (iv) a rare C3 type phage, isolated from human wastewater, overcomes this inhibitory activity due to an enhanced ability of the phage to bind to heparan-sulfated proteoglycans present in mucus or immobilized on the surface of intestinal epithelial cells, which likely drives the positional targeting of the phage to the exact ecological niche as the host bacterium. In addition, our data reveal that this treatment when compared to antibiotic treatment, did not alter the intestinal diversity of the microbiota. Taken together these data suggest a new mechanism of predation by a phage, one which may be highly useful for the killing of bacteria in intestinal environments and biofilms, both of which would be desirable properties of a targeted, specific antibacterial.

Potential narrow or limited ecological range of bacteriophages has not been as explored as much as their narrow host range capabilities. Using an experimental process of elimination, we determined that mucin can greatly inhibit phage infection. Mucins are composed of tandem repeats of serine and threonine that act as an attachment sites for o-linked glycans (N-acetlygalactosamine, N-acetylglucosamine, fucose, galactose and sialic acid) (McGuckin *et al*., 2011; Wang and Hasnain, 2017). Gel-forming mucins, which make up the mucus intestinal layer are large polymers (up to 40 MDa) of mucins attached via disulfide linkages (McGuckin *et al*., 2011). When we incubated phages with porcine gastric mucin, composed of the gel-forming mucin MUC 5AC (Sturmer *et al*., 2018), we saw reduced or no bacterial killing. However, when we added N-acetyl cysteine (NAC), which is known to exert a mucolytic effect by reducing the disulfide bonds that keep these mucin polymers together, bacterial killing was restored (Aruoma *et al*., 1989; Sadowska *et al*., 2006). NAC is a drug has been tested and used in the clinical setting to treat syndromes including cystic fibrosis (CF) and chronic obstructive pulmonary disease (COPD). In both of these cases it loosens thick mucus in lungs, but it can also serve as a treatment for acetaminophen overdose due to its antioxidant action (Sadowska *et al*., 2006; Saito, Zwingmann and Jaeschke, 2010; Bear, 2013). Perhaps this drug could also be used as an adjuvant to phages to help treat patients with bacterial infections in mucin-rich ecosystems such as the gut or the lungs of CF patients. We tested oral NAC treatment in mice and found it to be more effective in improving phage-mediated bacterial killing in the small intestine, but not in the large intestine. NAC has been shown to be rapidly absorbed following oral dosing after only 60 minutes (Tsikas *et al*., 1998). An oral dose of NAC has a short half-life of 2.5 hours and 10% bioavailability in the gut (Tsikas *et al*., 1998). It is likely that NAC did not exert a mucolytic effect on the distal large intestine. Interestingly, we observed elevated ExPEC levels in intestinal tissue with NAC treatment. It is possible that some pathogens like ExPEC may thrive in environments with a reduced mucus layer.

Another possibility, distinct from adjuvating phage with NAC, would be to search for phages that have evolved phenotypes that enhance their ability to encounter their bacterial host in a specific ecosystem, such as the GIT. Phage ES17, originally isolated from human sewage, is a member of the family *Podoviridae* and has a rare C3 type elongated capsid morphology with paddle-like short tail fibers (Ackermann, 2001). Previously, this phage was shown to be similar to PhiEco32, a rare Kuravirus which are known for their prolate head (Gibson *et al*., 2019). Phage ES17 showed a significant killing effect in cecal medium (CM) and in a mouse model of intestinal colonization compared to phage HP3. Phage ES17 bound mucin significantly better and coating bacteria with mucin increased phage adsorption. There is precedence for this concept. The bacteriophage adhering to mucus (BAM) model proposes that phages bind to mucus via *hoc* proteins or Ig (immunoglobulin fold-like) domains present on the capsid proteins of phages similar to T4 (Fraser *et al*., 2006; Barr *et al*., 2013, 2015). This mechanism was suggested to explain how phages bind to the metazoan mucosal surface and increase the probability of encountering a bacterial host, likely via a unique type of controlled diffusion that increases spurious interactions with its bacterial target (Barr *et al*., 2013, 2015). These studies showed that T4 adsorption increased in the presence of mucin from 65% to 80% absorbed phage within 10 minutes (Barr *et al*., 2015). The adsorption changes in phage ES17 in the presence of mucin were all or nothing from 0% to 98% within 10 minutes. This suggests that mucin may be a type of “bridge receptor” for phage ES17 since this phage is ineffective at killing in the absence of mucin, even at very high levels of phage. Such a model would make sense in the context of *E. coli* embedded in a mucin matrix, perhaps breaking down mucin as a source of carbon and thus binding it, thereby allowing the phage to adapt a capsular polysaccharide binding mechanism to structurally related sugars that are prominent on proteoglycans.

Bacteriophages have been shown to have a variety of carbohydrate-binding proteins present on tail fibers in order to bind different sugars present on the bacterial cell surface (Nobrega *et al*., 2018). Some of the most diverse are capsular depolymerases, which bind and break down capsular carbohydrates secreted by bacteria (Pires *et al*., 2016). These capsular depolymerases are diverse because of the diversity of bacterial capsule types. Capsule types have been shown to mimic some components of the intestinal system, including sugars present in the mucus layer (Vimr and Lichtensteiger, 2002; Severi, Hood and Thomas, 2007). Bacteria can use these mechanisms to subvert the immune system and to allow for the colonization of hosts (Vimr and Lichtensteiger, 2002; Kahya, Andrew and Yesilkaya, 2017). Phage ES17 was effective at killing bacteria in biofilms and possessed a protein with putative capsular depolymerase domains in a tail fiber protein. However, we could not detect any polysaccharide lyase activity and instead detected strong and stable affinity for a human intestinal sugar, the glycosaminoglycan sugar, heparan sulfate. This suggests a model whereby phage ES17 adapted to “colonize” the mucosal environment by binding to mammalian sugars. Heparan sulfate proteoglycans (HSPG) are expressed on the surface of many types of cells and prominently on the surface of intestinal epithelial cells (Garcia *et al*., 2016). One of the more tantalizing discoveries here was that human enteroids derived from colonic stem cells specifically bound phage ES17, but not other *E. coli* phages. HSPGs consist of repeating units of sulfated polysaccharides—heparan sulfate (HS). HS is a linear polysaccharide that begins as the precursor, heparan (disaccharide of *α*1,4-linked N-acetylglucosamine (GlcNAc) and *β*1,4-linked glucuronic acid (GlcUA)) (Sugahara and Kitagawa, 2002; Murphy *et al*., 2004). Subsequent modifications due to sulfation and epimerization leads to the mature form polysaccharide HS (Murphy *et al*., 2004). Bacteriophage K5 utilizes a tail spike lyase (*KflA*) to bind and degrade K5 capsule present on *E. coli* strains (Hanfling *et al*., 1996). Because the K5 capsule (heparoson; *β*1,4-linked GlcUA and *α*1,4-linked GlcNAc) polysaccharide has been shown to be structurally identical to heparan sulfate precursor, heparan, this enzyme can also act a heparinase (Hanfling *et al*., 1996; Roberts, 1996; Murphy *et al*., 2004; O’Leary, Xu and Liu, 2013). K5 heparan lyase cleaves the linkage of N-acetyl glucosamine and glucuronic acid, but is inhibited by sulfated regions (Murphy *et al*., 2004; O’Leary, Xu and Liu, 2013). Further homology searches showed that the carbohydrate binding domain found in ES17-TFP is structurally homologous to the binding domain present in the tail spike in phage K5. Considering that heparanase III cleaves at the same regions where K5 lyase would bind and degrades suggests ES17-TFP is utilizing the same receptor. Indeed, the addition of heparanse III, from the soil bacteria *Flavobacterium heparinum*, which specifically cleaves unmodified (unsulfated) NAc domains (GlcNAc-GlcUA) and NA/NS domains (GlcNS (N-Sulfo-D-Glucosamine) and GlcNAc) of HS, leads to abrogated phage ES17 binding to cells, meaning ES17 was highly specific for this type of glycan (Murphy *et al*., 2004; Park *et al*., 2017). Structurally similar mucins to heparan sulfate (Class III type, *α*-linked GlcNAc) suggest that this protein may also target these intestinal mucins (Ota *et al*., 1998; Fujita *et al*., 2011). This interaction is driven by phage ES17’s tail fiber protein, which is not related to the *hoc* proteins or contain an Ig-like fold as predicted from the BAM model. Thus, it would seem that phage ES17 uses a novel mechanism to localize not only a mucin-rich surface, but also directly to the mammalian epithelial surface (Figure 7). It is worth speculating that not only would this be a way to intercept *E. coli* that would also colonize this surface, but it could also be exploited therapeutically as a prophylactic “epithelium protecting” coating that prevents the pathobiont from invading the mucosa. Several pathogens, including bacteria and human pathogenic viruses have been shown to bind to HS and use them as receptors (Alvarez-Dominguez *et al*., 1997; Fleckenstein, Holland and Hasty, 2002; Cagno *et al*., 2019). It is tempting to speculate that this property might have been also acquired by phage, as suggested here.

**Figure 7:**
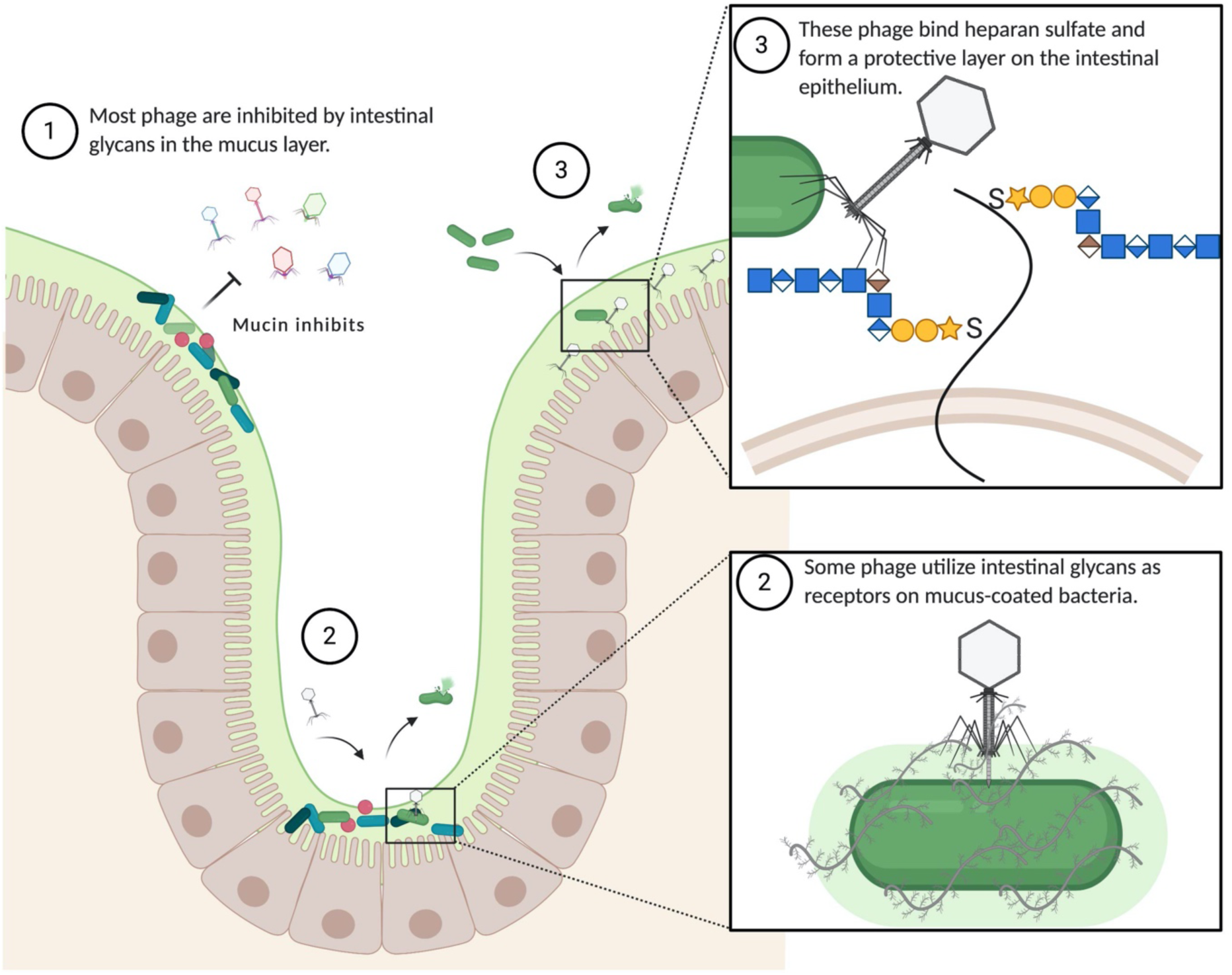
Model showing (1) mucins from the intestinal mucus layer inhibit phage infection, (2) phage ES17 can bind to mucin and utilize other intestinal glycans as a receptor to infect and kill mucus-coated bacteria, and (3) phages like ES17 can be utilized to coat the intestinal epithelium by binding heparan sulfate glycans to protect from invasive pathogen infection. Figure created with BioRender.com.

## Materials and Methods

### Bacterial strains and phages

ExPEC ST131 isolate JJ1901 was used in all ExPEC infections, except Figure S1Aiii (JJ2528). Both isolates were previously obtained from Dr. Jim Johnson (University of Minnesota)(Green *et al*., 2017b). Commensal *E. coli* ECN (Figure S1iv) was isolated from a human fecal sample. Deidentified strain DS515 is a clinical isolate obtained from the clinical microbiology laboratory at the Houston Veterans Administration Hospital. Prior to infections all strains were grown overnight at 37°C from a single colony streaked on an LB agar plate. The intestinal pathogen EAEC 042 (serotype 44:H18), was originally isolated from a child in Peru, and extensively characterized (Nataro *et al*., 1985).

Phages HP3, ES12, ES17, ES19, ES21 and ES26 have been previously described and characterized (Green *et al*., 2017a; Gibson *et al*., 2019). Phage 6914, 6915 and 6939 were recently isolated from sewage. All phages described in Table 1 were isolated by single plaque isolation from environmental sources as described previously (Green *et al*., 2017b; Gibson *et al*., 2019).

### Murine infections

Mixed ages (6-10 months) and sexes of BALB/c (Jackson laboratories, Bar Harbor, ME) mice were used in mouse models of infection. Mice were kept in SPF environment at Baylor College of Medicine CCM (Center for Comparative Medicine) Taub facility. All methods performed on mice were approved in accordance with relevant guidelines and regulations from “The Guide and Care and Use of Laboratory Animals” (National Institute of Health) and approved by Baylor College of Medicine’s Institutional Animal Care and Use Committee (protocol AN-6372). For infections mice were kept in a biohazard facility with sterile food and water. The mice were individually housed during colonization experiments and bedding replaced with autoclaved techboard liners for daily fecal collection. For colonization experiments, sample size was determined based on previous colonization experiments in mice (Green *et al*., 2017a). Mice received a 10^9^ CFU dose of ExPEC strain JJ1901 via oral gavage. Rodent health was monitored daily for indication of pain or disease. Bacterial colonization (fecal and intestinal) was determined after homogenization and selective plating for the chloramphenicol resistant strain JJ1901 on LB agar plates containing chloramphenicol and colony counting.

Purified phage in 3% (m/v) NaHCO₃ was administered either via gavage or in water 5% (m/v) sucrose added *ad libitum*. All groups received sucrose and NaHCO3 in water for consistency. The antibiotic ampicillin (1g/500 ml) was administered in water. Phage colonization was quantified after dilution of homogenates and serial plating on a double agar overlay assay.

### *Ex vivo* cecal model

A modified cecal assay was used for experiments (Theriot *et al*., 2014). Briefly, cecal contents from just euthanized mice were pooled and homogenized in sterile 0.09% NaCl solution at a 1:5 dilution (mg:mls). The homogenate was centrifuged to remove large particulates (2000 G for 30 sec.). The supernatant fluid was used for 4.5 hr phage killing assays at an MOI of 10 at 37°C, shaking (255 RPM), as previously described (Ma *et al*., 2018). All cecal and mucin experiments were performed using independent bacterial cultures grown up from different colonies streaked on a plate. This was considered a biological replicate. For FS CM, cecal supernatant was centrifuged (6000 G for 5 min.) and filtered through a 0.22 μm syringe filter. For HT CM, the supernatant was heated at 100°F for 20 min in a hot water bath, then cooled to RT for infections.

Insoluble CM and soluble CM (supernatant) were isolated post high-speed centrifugation (9000 G for 5 min) of CM. The insoluble pellet was resuspended in sterile 0.09% NaCl solution for infections (IN CM). For the mucin assays, porcine gastric mucin type II (PGM; Sigma-Aldrich) was used at various concentrations diluted in phosphate buffered saline (PBS). The mucolytic drug N-acetyl cysteine (NAC, Sigma-Aldrich, 5 mg/ml) was diluted in PBS for de-mucolytic assays.

### Phage Sequencing and Annotation

The annotation figure was generated from previous data (Gibson *et al*., 2019) using Geneious Prime 2019.2.3 and adjusted in Inkscape.

### Mucin-coating Adsorption and Imaging

For the adsorption curves, assays were performed at an MOI of 1 using mid-log phase cultures independently grown from different colonies on a plate (this was considered a biological replicate) and samples taken every 5 min for 10 min. Prior to adsorption, PGM (0%, 0.5% and 1.5% m/v) was added to the bacterial cultures for 10 min. RT, shaking (255 RPM). The cultures were centrifuged (6000 G for 5 min) and gently washed with PBS. The adsorption rate constants (K) were determined from the natural log of the slope of the adsorption curve versus the bacterial concentration. Some cultures were fixed in glutaraldehyde solution after 10 min adsorption for TEM imaging. TEM imaging was performed at the Texas Children’s Hospital Center for Digestive Diseases. After fixation the samples were dehydrated in ethanol and followed by evaporation of ethanol and coating with gold for observation during TEM.

### Mucin Binding ELISA Assay

Clear-walled Immulon 2 HB 96-well microtiter plate (Immunochemistry Technologies #227) were used for the ELISA assays. PGM (200 μl of 1mg/ml) was added to a microtiter plate and incubated at 4°C overnight. The next day the mucin was removed and wells were washed twice with PBS. The phage was added to wells for 1 hr then washed three times in PBST (PBS with 0.1% Tween-20). The wells were blocked with BSA then incubated with antibodies for the phage overnight at 4°C. Following washing steps, a HRP (horse radish peroxidase) conjugated antibody was added for 1 hr. To assess phage binding, TMB solution was added until the wells turned light blue and then a stop solution (2M H2SO4) was added. The absorbance was read at 450 nm. Each well was considered a biological replicate for this experiment.

### Biofilm Screening

Biofilm experiments were recently described but with some modifications (Mapes *et al*., 2016). *E. coli* DS515 cultures were independently grown from different colonies on an agar plate in tryptic soy broth (TSB) were diluted to an OD of 0.1 and seeded to inoculate a 96 well plate for the biofilm formation assay. Each well was considered a biological replicate. The plates were incubated statically at 37°C for 48 hrs. Following incubation, planktonic cells were removed, and the wells were washed three times in PBS. Phages diluted in TSB to 10^9^ PFU/ml were added and incubated for 24 hrs at 37°C statically. The wells were washed three times in PBS then an MTT assay was performed to determine the metabolic activity of the phage resistant biofilm-forming bacteria. Protein quantification was determined using a Bradford assay (Biorad) using BSA as a standard. Purified ES17-TFP was added at a concentration of 0.17 mg/ml to biofilms with added phage.

### HIEM Infection and Imaging

Human enteroid monolayers (HIEMs) were differentiated for 5 days (>90% confluent) as described (Poole, Rajan and Maresso, 2018). For experiments each well containing HIEMs was considered a biological replicate. HIEMs were incubated with phage at 10^8^ PFU/ml in culture differentiation media for 1 hour at 37°C in the presence of 5% CO2 in a humidified incubator, then washed in PBS. The HIEMs were fixed in Clark’s solution for 10 min to preserve the mucus layer. The HIEMs were permeabilized and blocked with 5% BSA in 0.1% Triton X-100 in PBS for 30 min at RT. Mucus was detected using antibodies to MUC2 (1:200) (Abcam) and nuclei stained with 4′, 6′-diamidino-2-phenylindole (DAPI) (300 nM) for 5 min at RT. Antibodies against phages HP3 and ES17 were generated from whole virus (phage) injection into rabbits performed by Pacific Immunology. A 13 week antibody production protocol consisted of 4 immunizations and anti-sera collection.

To selectively removed HSPG from GAG chains, enteroid cultures were pretreated with heparinase III (Sigma, 2U/ml) for 2 hours, as described (Jiao *et al*., 2007), followed by the addition of phage. Images were captured using a Zeiss LSM 510 confocal microscope. Represented images were adjusted equally for brightness and contrast using FIJI software version 2.0.0. The images were adjusted equally for brightness and contrast. Particle analysis was used to determine particles per well. The Cell counter program in FIJI was used to quantitate bacteria (EAEC) per field of view.

### EAEC infection and staining

For infection of HIEMs, the EAEC strain 042 was grown overnight and then sub-cultured (1:100) for 2 hrs at 37°C. One microliter of the log-phase bacterial culture was added to 100 microliters of the cell culture media and added to chambered slides either with phage-coated HIEMs (as described above) or untreated. Cultures were incubated for 3.5 hrs at 37°C in the presence of 5% CO2 in a humidified incubator. Following growth, bacteria were removed, by gently washing the cells in PBS then the remaining cells were fixed (Hema 3 fixative) and stained using a Geimsa-Wright stain (Fisher scientific). The infection and staining technique have been previously described (Rajan *et al*., 2018).

### Statistics

Statistical analysis was performed using PRISM 8 software. Microbiome 16S data was analyzed using ATIMA (Agile Toolkit for Incisive Microbial Analyses) as described in supplemental methods. For figures with log transformed data and groups >2 significance was determined using a one-way ANOVA analysis (Figures 1, 2, 3, 4, 5 and 6) or two way ANOVA when necessary (Figure 3E). For multiple comparison analysis, the secondary test, Tukey, was used. For non-transformed data a normality test (Shapiro-Wilk) was performed to determine normality before a one-way ANOVA analysis was performed. Significance was determined to be a p value less than or equal to 0.05. Unless otherwise stated all graphs show the mean and standard deviation.

## Acknowledgements

This work is supported in part by grant from US Veterans Affairs (VA I01-RX002595), Roderick D. MacDonald Research Fund at Baylor St. Luke’s Medical Center, the Mike Hogg Foundation, and Seed Funds from Baylor College of Medicine Seed Fund We would like to thank Dr. Melinda Engevik for general expertise and protocol help related to mucin work. We would also like to thank Hannah Johnson from the Integrated Microscopy Core at Baylor College of Medicine for technical assistance related to imaging.

## Author Contributions

S.G., C.L., X.Y., W.S., A.R., H.C. and X.S. performed experiments and analyzed data. J.C. performed bioinformatics studies. R.R., B.T., and H.K., contributed to design of studies and edited the manuscript. S.G. designed studies and wrote the manuscript. A.M. contributed to design and overall major goals of the study and edited the manuscript.

## Supplemental Figures, Methods and References

**Figure S1.**
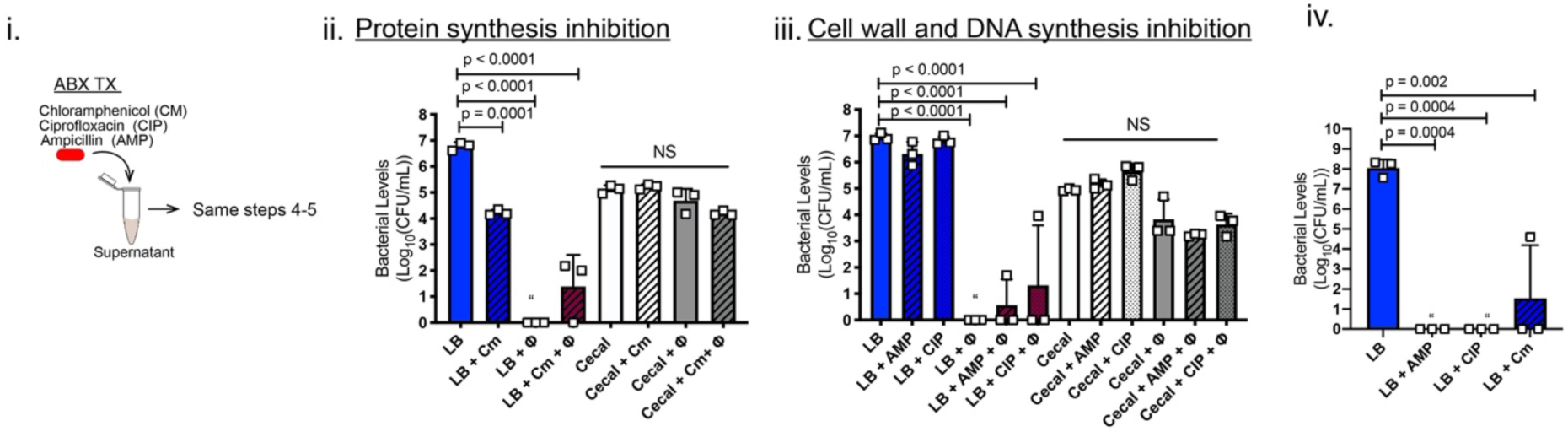
(i) ExPEC levels in CM assay incubated with antibiotics (ii) chloramphenicol (Cm) ciprofloxacin (CIP) or (iii) ampicillin (AMP). (iv) Commensal bacterial levels incubated in antibiotics indicated. (N=3-5). Exp. performed 1X. Squares indicate individual biological replicates from independent cultures. NS not significant.“ = none detected. Mean, ±SD shown.

**Figure S2.**
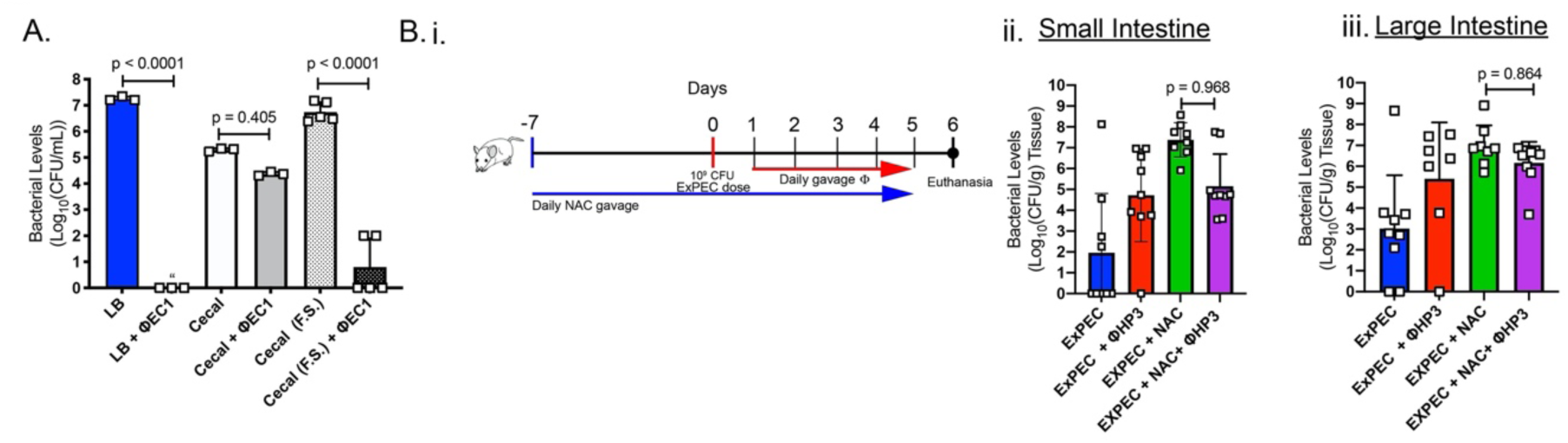
(A) ExPEC levels in CM (FS) with phage EC.1. (N=3) (Bi) Mice were gavaged daily with N-acetyl cysteine for 2 weeks. ExPEC was gavaged on day 0 following protocol from Figure 1A. (Bii) Small intestinal ExPEC levels. (Biii) Large intestinal ExPEC levels. (N=9). Exp. performed 1X. Squares indicate individual biological replicates from independent cultures or individual mice. NS not significant.“ = none detected. Mean, ±SD shown.

**Figure S3.**
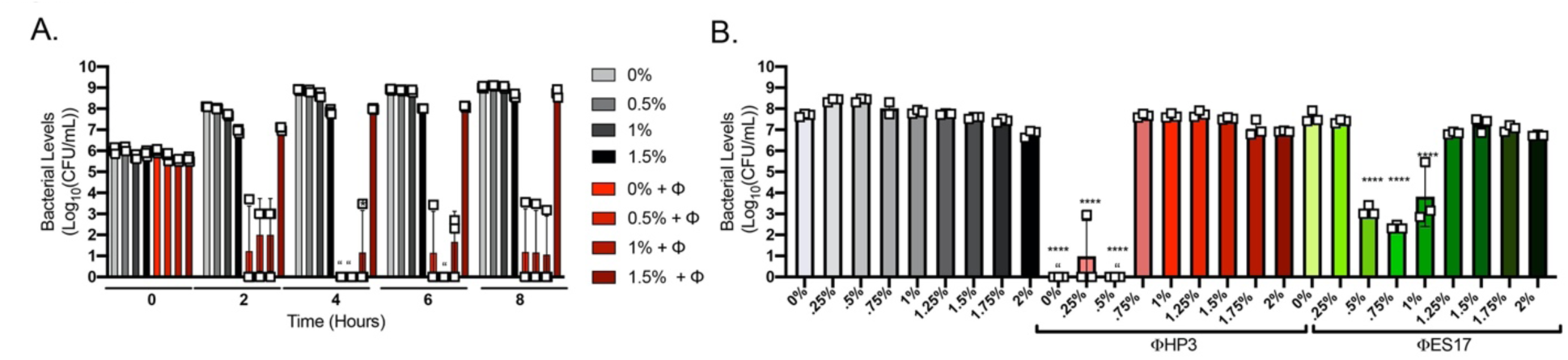
(A) ExPEC levels from 0 to 8 hours in LB plus mucin concentrations indicated and +/- phage HP3. (B) ExPEC levels with phages HP3 or ES17 (MOI of 10) in LB plus mucin concentrations indicated. (N=3). ****p < 0.0001. “ = none detected. Exp. performed 1X. Squares indicate individual biological replicates from independent cultures. Mean, ±SD shown.

**Figure S4.**
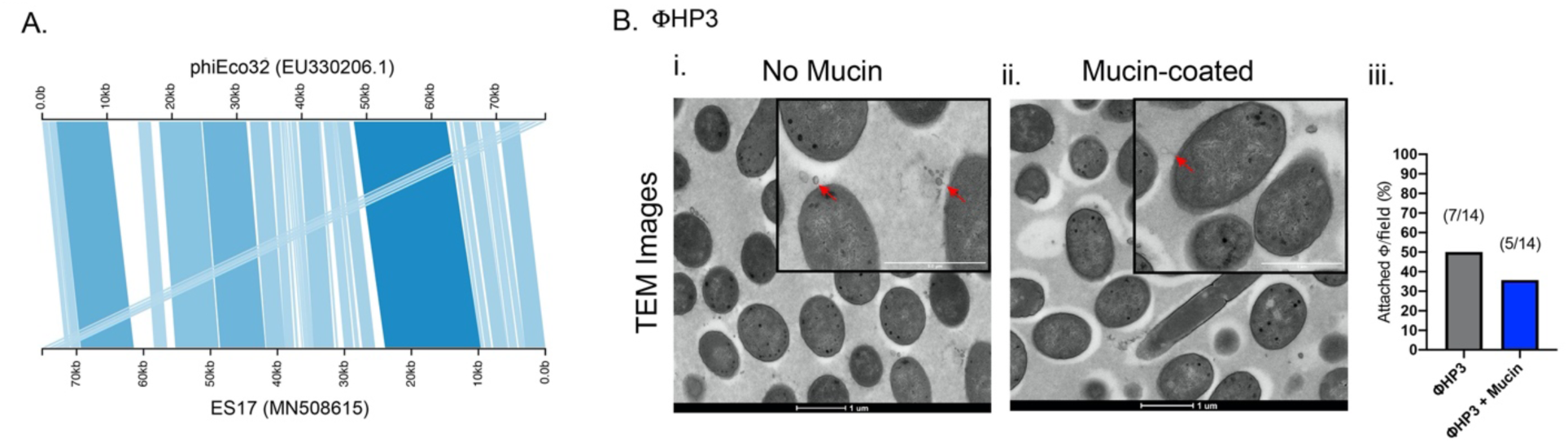
(A) Graphical representation of BLAST analysis of phage ES17 (bottom axis; MN508615) and phage phiECo32 (top axis; EU330206.1) using Kablammo web-based software (Wintersinger and Wasmuth, 2015). Trapezoids drawn between the axis indicate individual BLAST alignments between the two sequences. The stronger alignments are shaded darker. (Bi-ii) TEM images of phage HP3 on ExPEC fixed 10 min post-incubation. White bar scale is 1 μm. Red arrows indicate phage. (Bii) Quantification of attached phages per field view. Mean, ±SD shown.

**Figure S5.**
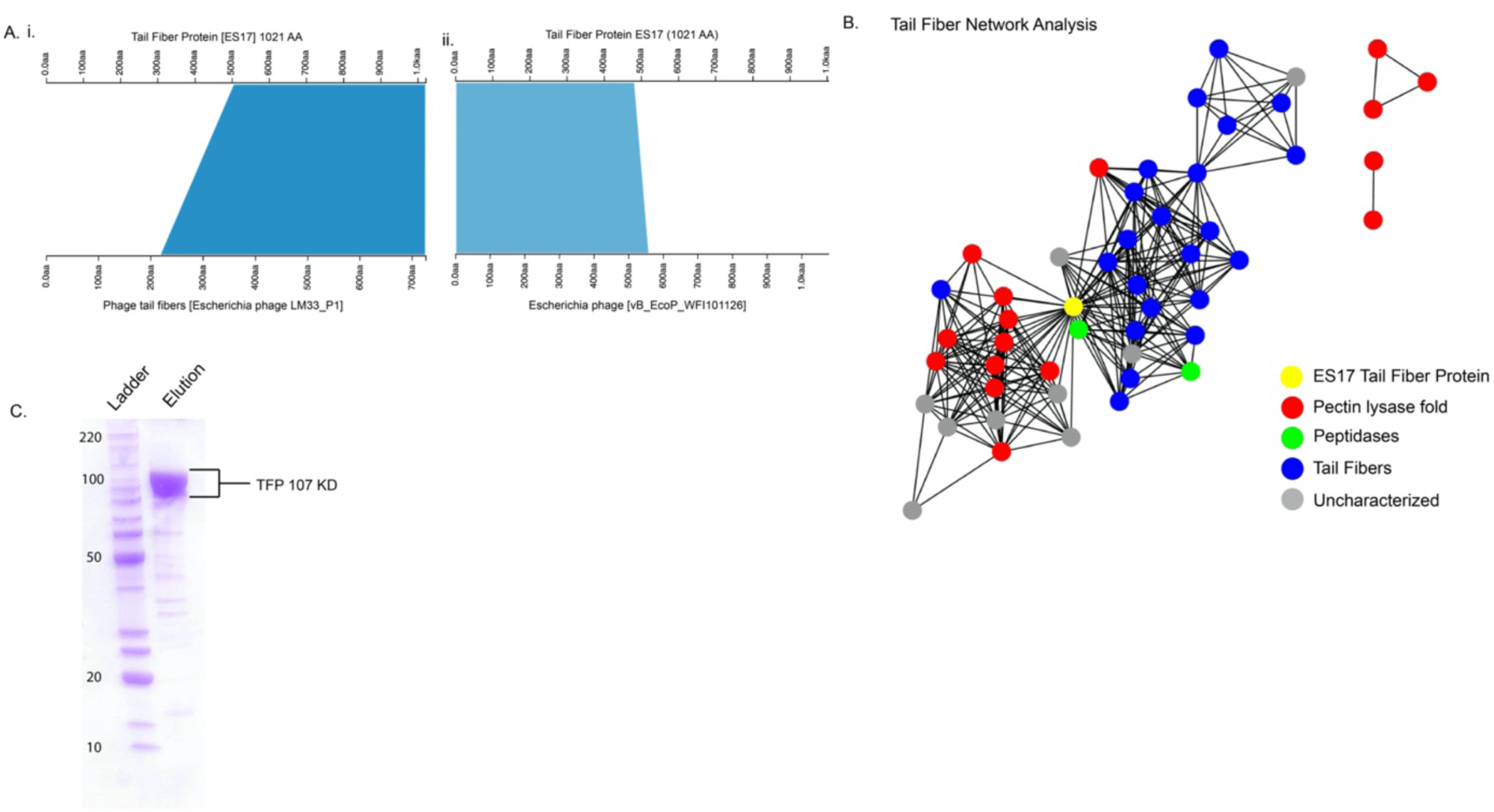
(Ai) BLAST analysis of ES17-TFP (top axis) and phage LM33_P1 tail fiber (bottom axis, ref|YP_009324518.1|)) and (Aii) *E. coli* phage vB_EcoP_WFI101126 tail fiber (bottom axis, gb|QBQ76440.1|) (B) Network pictured. An E-value of 1E-1 was used as a cut off for node inclusion. Each node represents a single protein. Edges connect nodes with a BLAST E-value of less than 1E-10. The yellow node is the ES17-TFP, red nodes are proteins that belong to the pectin lysase fold/virulence factor family (IP011050), green nodes are proteins that are predicted peptidases; blue nodes are phage tail fibers; and gray nodes are uncharacterized proteins. (C) Protein SDS-PAGE gel showing purified ES-TFP pooled elutions and protein ladder.

**Figure S6.**
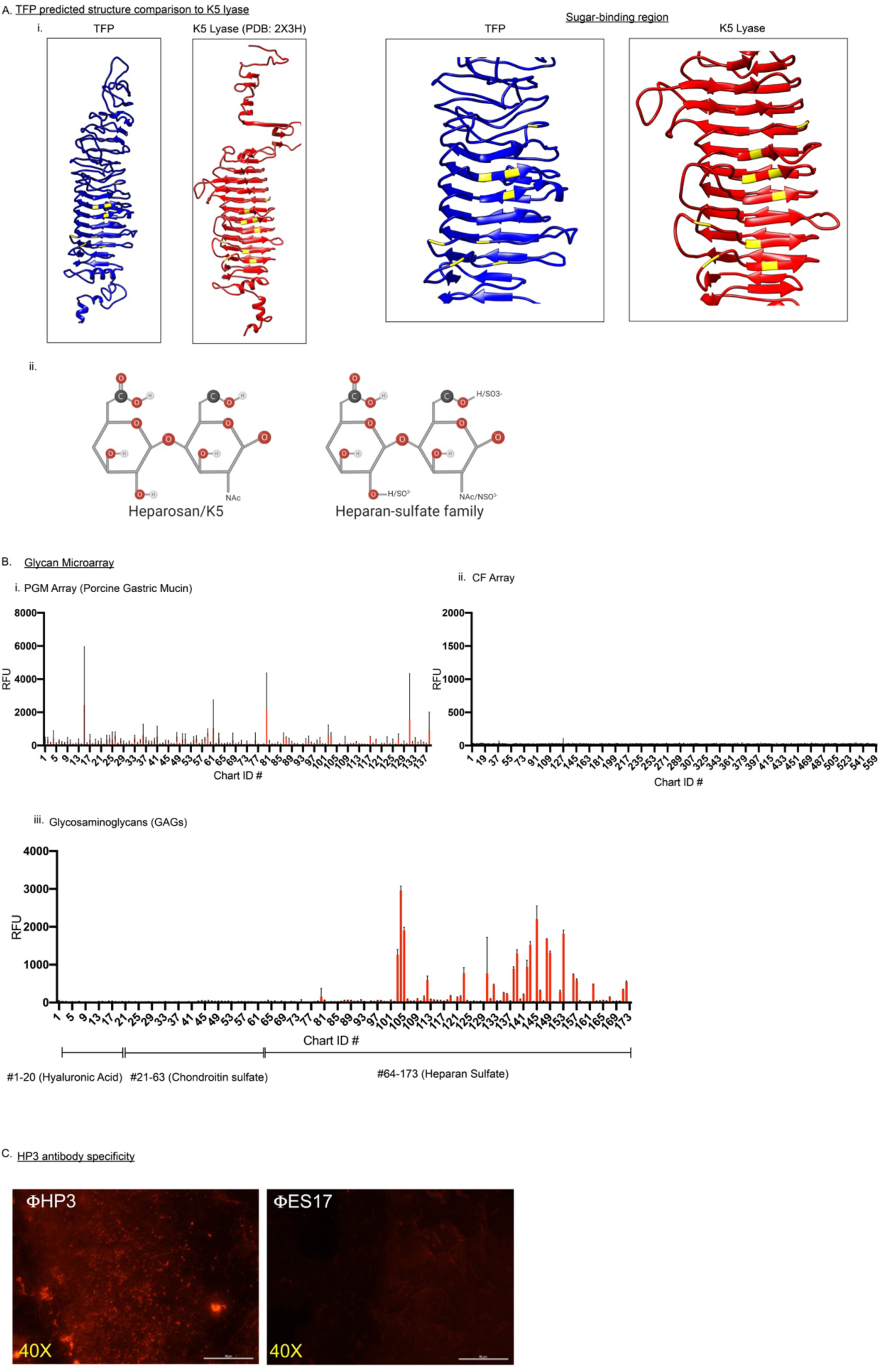
(Ai) Structures show the predicted structure of tail fiber (blue), the structure of K5 lyase (PDB:2X3H; red). Identical residues between K5 lyase and ES17 tail fiber protein are yellow. (Aii) Structures of heparosan or K5 capsule and heparan-sulfate derivative. (B) Glycan microarray analysis in relative fluorescent units (RFUs) using purified ES17-TFP as the GBP (glycan binding protein) and glycans derived from purified (Bi) porcine gastric mucin, (Bii) other mammalian glycans (CF array, see methods) and (Biii) glycosaminoglycans. N=3. Experiment was performed 3X. (C) Immunofluorescent staining of ES17 and HP3 phages fixed on a slide using antibodies generated against HP3 and secondary fluorescently conjugated (Alexa Fluor 594; Red).

## Supplemental Methods

### Antibiotic treated CM

For antibiotic treated cecal medium, CM was incubated with chloramphenicol (10 ug/mL, EMD Millipore) or ampicillin sodium salt (100 ug/mL) or ciprofloxacin (10 ug/mL) throughout the 4.5 hour infection using ExPEC JJ1901 (Figure S1ii) or JJ2528 (Figure S1iii) or commensal *E. coli* ECN (Figure S1iv).

### NAC Mouse Model

Mice (BALB/c) were housed individually and gavaged daily with NAC (N-acetyl cysteine) at a concentration of 40 mg/mouse/day throughout the experiment as indicated (14 days). This was a tolerable concentration and shown to be effective in the mouse small intestine (De Lisle, Roach and Jansson, 2007). Mice were infected as described in the primary methods, then euthanized on day 6 and plated to determine bacterial concentration in the intestinal contents.

### Genome Comparison

Genome comparison was performed using a BLAST analysis of phage ES17 (MN508615) and phiECO32 (EU330206.1) and visualized using Kablammo software (Wintersinger and Wasmuth, 2015).

### ES17-TFP Construct Generation

The construct used to purify tail fiber protein (TFP) was prepared by Genewiz (South Plainfield, NJ) using established protocols. The construct sequence for TFP was synthesized from the phage ES17 sequence (MN508615.1) nucleotides 58,576-61,641, ttaacgcagggcacgtctactattagcgaagcgcatagcctctgcctgagtagtgaactccgctacgaagtaatcagcga ggcgtaggactactccgcctgtaccggctgtacttaggcgcaggttcactgaaccggaagcactagcctgagctacacca aaggtactcacccactggccccccttagtaaatacttcacctacattgatatctccaccgatgcggaagtttgtaggggttcc ctgtaccagttcagtgctaatgccccatacataccagttaccggaagttatctgaaatgcagcaccagtaagctcatactca gcagcaccagggatttcgaagtacatgcagctctgagataactggccttgtggctgcaaactcccagtgctgcttgccgta ccactaaattcagcagtacctccatcgaagggtttaactaccagcttattaccaggtacacgatagatgtatccttccggca ggtatccacgcatactcatattaggtgtccagctttgcccaatcttgcctgtctgagccgctacagaacgtaccattacatcttt gccggagctaccattcaggtacaattgatcagccacaaaactgctgttggcatctacagagatatccatgtggccatctga attatctactcggcagccataagtaacaaggttggagttctgaagctctacgttattaaggtatgtgtactctgcaaagaagat actggtatcaataagtttaaggacacgaggcttctgagatacaccatctacattaatagctggagtagtagcgatagcttcc atccatacattggatacaccacatggggcagtagggcagtcaccatccttgttcttgaggtaaataccaccaccgccaga ggcttccataatactatctttaatccaccagccgcctacaccacctccgaatgtaccaggcgcttctcctgttgtagcgttcaa gtaaacacagtaagtgctaataccatcaaagtgcatgttctgaagggtgtcagagcctaagtgcatctcagaagcagact gtgcccagtaaccccagtcacaaccttgcacgccaatgcctttccatgtattacctatattcccggaaggtttctgaatggcct tgttaatgttatggatgtatacgtcactaaagttgtgacgaccagcatattggtcggtcgggtcaaaactaatacctacagcc ccggttgtcccagaaccatctatagcaaaaccaccactatggcggaattgccaatgaggcgtagatgccttaacctgaat agtgtctgtaaagtttgacggagctttcagtaccgtggctccaccaacaccctttaactgtgtactaccaatagtaatcgcttc actgtattcaccgggggaaacaattaagtcgttttttggtaacgatgcatttaccgcagcttgtaaggtagggaaatctcctga atgcttaatgtcgttgttacggacaagcaaacttctctcaccactgccaatgtttttagccccatctttactggaaagactctca acagaattaccattttcaaaggtaataagggatgcaccttcacctgcattactacttttcagccttgccagcaaatctgcttca cttgtatgcaactgtctcaaagcaaccgcgtcattatcatctacagcatcaggaattactggagattgtgtgaaagtaacctt gttggtcatcgaggtgatatcagagttatcaccactcttagctgcaactagagctagtcgagcattctctgatgaagtagctcc tgtaccacctttggtaataggtacaattgattcagtggaaaccaaacctaaaccaagtttagtacgtgcagcacttgcagta gttgcccctgtgccaccctgagagatagtcacagggcgagcggaagacagctcttctcgtactgtaaatactcgggaacc aactcctccagtccagcttacagagaaaactgagtagtagttattattgtttgtccctggcaccactcgtacctcacctcctcct gtaactacgccaatgacttcaagatataacccataattggcaggataatttatctcagcaggggcatttgtaatattcgacatt aatactgaatagttacctccattgacaaaattaaatgtttgccagtccatagccttcactttggttaaggtagggctaccaatcc catgctcagcgagtccaatatttgtaatagctgcacttgtgtcggttatttcggataggttattagttacactaaaagctcccact tgagcagctgtgtagtctccagtagcaggaactacagcacccttacgtccattgaatgtggagacagaatcaaaaaccac tgaacctgtttgttcagcgccgccattgattgaaacagatcgaacagatggaatcaagttggcaatatctgacatgtcagct cggtataaaccaccatcatctttatctacatagaaaccatagtcatcaacaacaggggtaccaacaggaacgcttccaat atcaccgattgttccgtcagtaccatcacgccccggtttaccttctggaataccaaaagcaaagacacctgttgcattatcat aagaccccgtagctaggcttcccggaggaagaagcgttgtttgaactgtcatgttagagataatgtcaagttgtgctagaac gttttgagttgcctcttttgaaagctcagttgctgcattggtgttgtcaattgcagttcctgtggattgaatcaaacctaagagcac ttgttcttgtgactcaaagtcactcttgagttgtttaatgtcttcttccagtagatgaccttgagcatacagacgttctacttcagcc aaaatctcctctactgaggagtatttagtttgggacaacaatgcccaatatttagagtctgctgcatattcagctgctgatttata agcaccaatagttccctcagtagctccaaattgtcctacattgttaacattcttaggtgcttggttgttataaacgatcat.

This sequence was ligated onto pET-21b (+) (ampicillin) vector using restriction enzymes 5’ *BamHI* and 3’ *Xhol* by recombination and delivered as a mini-scale DNA sample.

### Protein Purification

BL21 (DE3) cells were used for expression of His6-TFP under the control of a T7 promoter expression. Cells were induced with Isopropyl ß-D-1-thiogalactopyranoside (IPTG) 1mM for 2 hrs at 37°C. Then the culture was pelleted at 7000 RPM for 20 min and resuspended in Binding buffer (50mM NaH2PO4, pH 8.0 0.5 M NaCl). Cells were lysed using a French Press. Following lysing and centrifugation (15,000 RPM for 20 min.) the lysate was added to a Ni-NTA column for binding, washing 4 times in buffer (Binding buffer with 20mM imidazole) and elution (binding buffer with 250 mM imidazole). Buffer exchange was performed in PBS. An SDS-PAGE gel was run to verify the size of the protein along with a ladder (Benchmark Protein Ladder).

### HIEM Generation

Human Intestinal Enteroid (HIEs) were cultured from 3D enteroid cultures derived from human colonic biopsy tissue using methods developed by Sato and Cleavers and 2-D enteroid monolayers (HEIMs) developed using methods of Van Dussen et al. (Sato *et al*., 2009, 2011; VanDussen *et al*., 2015; Saxena *et al*., 2016). Briefly, HIEs were derived from human biopsy samples. The colonic crypts were isolated, embedded in Matrigel (Corning) or collagen (Sigma) and incubated with complete growth media with growth factors (CMGF+) for differentiation into 3D enteroid structures. Next, chambered slides (Greinier Bio-One) were coated with Matrigel or collagen, dissociated, about 50 spheroids, (3D enteroids cultured CMGF^+^ for 7 days) were added per well. HIEs were dissociated using 0.05% trypsin/0.5mM EDTA and a gravity flow strainer. Cells were allowed to grow and adhere 1-2 days with CMGF^-^. Then the media was removed and replaced with Differentiation media. Typically for experiment, HEIMs were used 4-7 days later when fully confluent (>90%) and differentiated. For our experiments this was day 5. The components of media the CMGF+, CMGF- and Differentiation media are described previously (Poole, Rajan and Maresso, 2018).

### Glycan Microarray

Glycan microarray analysis was performed by the Emory Comprehensive Glycomics Core using the methods described below.

An oligosaccharide array composed of 560 defined glycans from the Consortium for Functional Glycomics were obtained from MicroArray Core Facility of the Scripps Research Institute (La Jolla, CA) (lists can be found at http://www.functionalglycomics.org/static/consortium/resources/resourcecored2.shtml#Glycolipid) as described(Heimburg-Molinaro *et al*., 2011). The glycans were printed on Nexterion 3-D hydrogel coating (H) slides using a contact printer in replicates of six. A second microarray consisting of 138 glycans from purified porcine gastric mucin was also included. And a third microarray consisted of 170 charged glycosaminoglycans (GAG), including oligomers from hyaluronic acid, chondroitin sulfates and heparan, obtained from the Emory Comprehensive Glycomics Core (Atlanta, GA). Glycans were conjugated with the bifunctional linker (AEAB) and printed on Nexterion 3-D hydrogel coated (H) slides using a contact printer Aushon 2470 (Quanterix, Billerica, MA) in replicates of four. His-tagged TFP (up to 40 ug/mL) was diluted in PBS buffer with 1% BSA and 0.05% Tween 20 and incubated on the microarray slides for 1 hr at RT. Slides were washed with PBS buffer with 0.05% Tween 20 and PBS buffer. Alexa Flour 647 anti-His-tag mouse mAb (MBL International, Woburn, MA) was diluted to 5 µg/ml and incubated on the slides for 1 hr. After washing and drying, the slides were scanned with an InnoScan 1100 AL scanner and the data were processed using Mapix 8.2.5 software (Innopsys, Chicago, IL). Mean relative fluorescence units (RFU) from replicate spots were averaged after subtracting local median background, and the standard deviation (STDEV) were calculated and plotted as error bars in the histogram plot of the glycan array.

### Bioinformatics modeling

The phage genome was annotated using the RASTtk (Brettin *et al*., 2015). The Enzyme Function Initiative-Enzyme Similarity Tool (EFI-EST) was used to visualize BLAST results of the tail fiber protein (TFP) (Gerlt *et al*., 2015). The network visualization showed that TFP was related to other tail fiber proteins and proteins that belong to the pectin lyase fold/virulence factor family of proteins (IPR011050). HHpred was used to identify other proteins with similar structures (Hildebrand *et al*., 2009). The highest scoring template was a Tailspike protein from *E. coli* bacteriophage HK620 (PDB: 2VJJ; E-value = 9.4E-23). The other top five hits from Enterobacteria or *Escherichia* phages were a tailspike protein from Enterobacteria phage SF6 (PDB: 2VBK; E-value=1.9E-14), a putative endo-glycosidase tailspike protein (PDB: 6NW9; E-value=3.1E-14), a colanidase tailspike protein from Enterobacteria phage phi92 (PDB:6E0V; E-value=1.2E-13), a sugar-binding tailspike from *Salmonella* phage Det7 (PDB:6F7D; E-value=2.1E-12), and a K5 lyase (E.C.4.2.2.7) tailspike protein from Enterobacteria phage K5 (PDB: 2X3H; E-value=4E-12). All of these tailspike proteins bind and degrade polysaccharides and are classified as members of the pectin lyase fold/virulence factor family (IPR011050)(Freiberg *et al*., 2003; Barbirz *et al*., 2008; Schwarzer *et al*., 2012; Broeker *et al*., 2019; Greenfield *et al*., 2019).The K5 lyase has well-known heparanase activity due to the similarities between K5 and heparan-sulfate (O’Leary, Xu and Liu, 2013). All six were used to produce a predicted structure of the ES17 tail fiber protein in Modeler (Zimmermann *et al*., 2018). Structures were aligned using UCSF Chimera. Chimera’s matchmaker function was used to structurally align the predicted tail fiber 2 protein with the HK620 tailspike and the K5 lyase. Solvent exposed identical residues between ES17 tail fiber protein 2 and K5 lyase were determined using Chimera’s match-align to determine what residues were both identical and within 5 angstroms of each other in the structural alignment.

### 16S rRNA Gene Sequencing

The Center for Metagenomics and Microbiome Research (CMMR) at Baylor College of Medicine performed 16S rRNA sequencing and analysis.

The 16S rRNA gene sequencing methods were adapted from the methods developed for the Earth Microbiome Project (Caporaso *et al*., 2012) and NIH-Human Microbiome Project (Caporaso *et al*., 2012; Huttenhower *et al*., 2012). Briefly, bacterial genomic DNA was extracted using the Qiagen MagAttract PowerSoil DNA Kit (formerly sold by MO BIO as PowerMag Soil DNA Isolation Kit). The 16S rDNA V4 region was amplified by PCR using primers 515F and 806R containing Illumina adapters and a single-index barcodes (Caporaso *et al*., 2012). The amplicons were visualized with gel electrophoresis and purified to remove primer-dimers and non-specific amplicons. The samples were quantified using the Qubit® fluorometer and pooled at a DNA mass of 100 ng per sample. The amplicon pool was sequenced on the Illumina MiSeq platform using the reagent kit v2 (2 × 250 bp) and the paired-end protocol yielding paired-end reads that overlaped almost completely.

The raw data files in Binary Base Call (BCL) format created by the MiSeq run were first converted into the FASTQ format and demultiplexed based on the single-index barcodes using the Illumina BCL2FASTQ software. The demultiplexed FASTQ read pairs were then merged using USEARCH v7.0.1090 (Edgar, 2010) ‘fastq_mergepairs’ function, requiring read pairs to overlap by at least 50 bp, a merged length of at least 252 bp, a truncation quality above 5, and zero differences in the overlapping region. The merged files were then filtered further, using the USEARCH70 ‘fastq_filter’ program, and only allowing for a maximum expected error of 0.05. Merged reads were then combined into a single FASTQ file, which was filtered for PhiX using Bowtie2 v.2.3.4.3 (Langmead and Salzberg, 2012) and the ‘very-sensitive’ parameter setting. After removing PhiX reads, the FASTQ file was transformed into a FASTA format file. The FASTA file was run through USEARCH70 ‘derep_fulllength’ program and the reads were sorted by size using usearch70 ‘sortbysize’ program. The reads were clustered into operational taxonomic units (OTUs) at a similarity cutoff value of 97% using the UPARSE algorithm (Edgar, 2013). This was accomplished in an iterative stepwise manner in increments of 0.4% using USEARCH70 ‘cluster_otus’ function. The output from the previous increment was fed into the next iteration, until a maximum of 3.2% was reached. All the intermediate files of this step were filtered for any chimeras and after the last run through the above loop, the final output was run through USEARCH70 ‘uchime_ref’ program against the GOLD database (Kyrpides, 1999; Mukherjee *et al*., 2019), using only the plus strand and allowing for no chimeras, in order to create a clustered OTU file with no chimeras. The OTU file was mapped against an optimized version of the latest current SILVA Database (Quast *et al*., 2013) containing only sequences from the V4 region of the 16S rRNA gene to determine taxonomies. This step was performed using USEARCH70 ‘usearch_global’ function, specifying the identity threshold to 96.8%. Abundances were recovered by mapping the demultiplexed reads to the OUT file and all files created in the loop were then run through a program developed in-house that resolved the iterative UPARSE steps, creating an OTU table in BIOM format and removing the chimera and singleton reads. The BIOM file was summarized, recording the number of reads per sample, and merged with a file that was generated for the overall read statistics, to produce a final summary file with read statistics and taxonomy information. ATIMA (Agile Toolkit for Incisive Microbial Analyses) was used to generate beta diversity graphs.

## Notes

### Competing Interest Statement

The authors have declared no competing interest.

